# KAT3 Shuttling Between Neuronal Identity and Activity-Dependent Plasticity Programs Drives Large-Scale Chromatin Remodeling

**DOI:** 10.64898/2026.08.28.747502

**Authors:** Sergio Niñerola, Beatriz del Blanco, Mirjam Cangonja, Juan Paraíso-Luna, Federico Miozzo, Patricia Torres-Raves, Michal Lipinski, José M. Santos-Pereira, Angel Barco

## Abstract

Activity-dependent transcription is a central feature of neuronal plasticity. Here, we show that neuronal activation triggers genome-wide redistribution of CBP and p300 in hippocampal neurons. Upon stimulation, KAT3 cofactors relocate from super-enhancers supporting neuronal identity to enhancers associated with activity-regulated genes, accompanied by transient changes in H3K27ac, chromatin accessibility, and three-dimensional genome architecture. Mechanistically, distinct TF families control KAT3 shuttling: proneural bHLH factors such as NeuroD2 maintain cofactor occupancy at identity-associated regulatory elements, whereas AP-1 binds de novo at plasticity-associated loci. This dynamic redistribution reshapes enhancer landscapes and chromatin interactions, enabling robust activation of plasticity genes while transiently attenuating neuronal identity programs. Remarkably, FOS overexpression is sufficient to reproduce the repression of neuronal identity genes observed during stimulation. Together, our findings reveal a reversible competition between transcriptional networks governing neuronal identity and plasticity and identify KAT3 redistribution as a key mechanism coupling neuronal activity to large-scale chromatin remodeling.

## Introduction

Activity-dependent transcriptional changes are a critical component of neuronal responses to stimulation and play a central role in information processing. They are initiated by increases in intracellular Ca^2+^ at activated synapses, which trigger signaling cascades that include the activation of multiple kinases and transcription factors (TFs) (Yap and Greenberg, 2018; Lee and Fields, 2021; Sugo et al., 2025). These TFs drive gene expression programs that prolong the effects of neuronal activation by inducing plasticity-related genes and promoting lasting changes in chromatin organization (Benito and Barco, 2015; Griffith et al., 2024). Notably, many of the same factors also play essential roles during development, when distinct combinations of TFs occupy cis-regulatory elements (cis-REs) to specify somatic cell fates and establish the transcriptional networks that define neuronal function (Heinz et al., 2010).

A key function of these TFs is to recruit transcriptional co-activators, such as the type 3 lysine acetyltransferases (KAT3) CBP (*aka* KAT3A) and p300 (*aka* KAT3B) (Malik et al., 2014). These co-activators are present in limiting availability in the nucleus (Goodman and Smolik, 2000; Dyson and Wright, 2016) and acetylate lysine residues on both histone and non-histone proteins while also acting as molecular scaffolds within transcriptional complexes that respond to synaptic input, thereby promoting gene expression (Lipinski et al., 2019). Although both KAT3 paralogs acetylate hundreds of lysine residues, their depletion has particularly strong effects on specific sites, including histone H3 lysine 27 acetylation (H3K27ac), a histone post-translational modification widely used as a marker of active enhancers (Creyghton et al., 2010) and particularly sensitive to CBP/p300 activity (Raisner et al., 2018; Weinert et al., 2018; Lipinski et al., 2020, 2022). Dysregulation of this pathway has been linked to memory impairments and neurological disorders, most notably to the intellectual disability associated with Rubinstein– Taybi syndrome (Alarcon et al., 2004; Korzus et al., 2004; Lopez-Atalaya et al., 2014), caused by pathogenic variants in *CREBBP* or *EP300*, as well as Alzheimer’s and Huntington’s diseases and age-related cognitive decline (Gräff and Tsai, 2013; Achour et al., 2015; Nativio et al., 2020; Signal et al., 2024).

Our recent multi-omic analysis of activity-driven changes in excitatory neurons *in vivo* during KA-induced status epilepticus (SE), a condition characterized by widespread neuronal activation, showed that the transcriptional burst induced by neuronal activity is accompanied by increased chromatin accessibility at activity-regulated genes and enhancers, as well as strengthened promoter-enhancer interactions (Fernandez-Albert et al., 2019). However, the epigenetic regulators that drive these changes remain incompletely defined. Given the broad role of CBP and p300 as coactivators and their essential functions in plasticity and cognition, they are strong candidates to couple neuronal activity to dynamic chromatin regulation. Here, we sought to determine how CBP and p300 are redistributed upon neuronal activation, define the chromatin changes associated with this redistribution, and uncover its underlying molecular mechanisms and transcriptional consequences. Our findings reveal a transient and reversible competition between transcriptional networks governing neuronal identity and plasticity, with potential implications for neuronal adaptability and dysfunction in neurological disease.

## Results

### Activity-dependent redistribution of KAT3 proteins

To investigate the role of CBP and p300 in activity-dependent chromatin remodeling during intense neuronal activation, we assessed their genome-wide distribution in the hippocampus under basal conditions and following SE using ChIP-seq. SE was triggered by intraperitoneal injection of kainic acid (KA; 25 mg/kg), and hippocampi were collected 1 h post-injection (**Fig. 1a**). ChIP-seq was performed using specific antibodies against CBP and p300. Principal component analysis (PCA) of the resulting datasets revealed a clear separation between conditions and coactivators (**Supp. Fig. S1a-b**).

**Figure 1.**
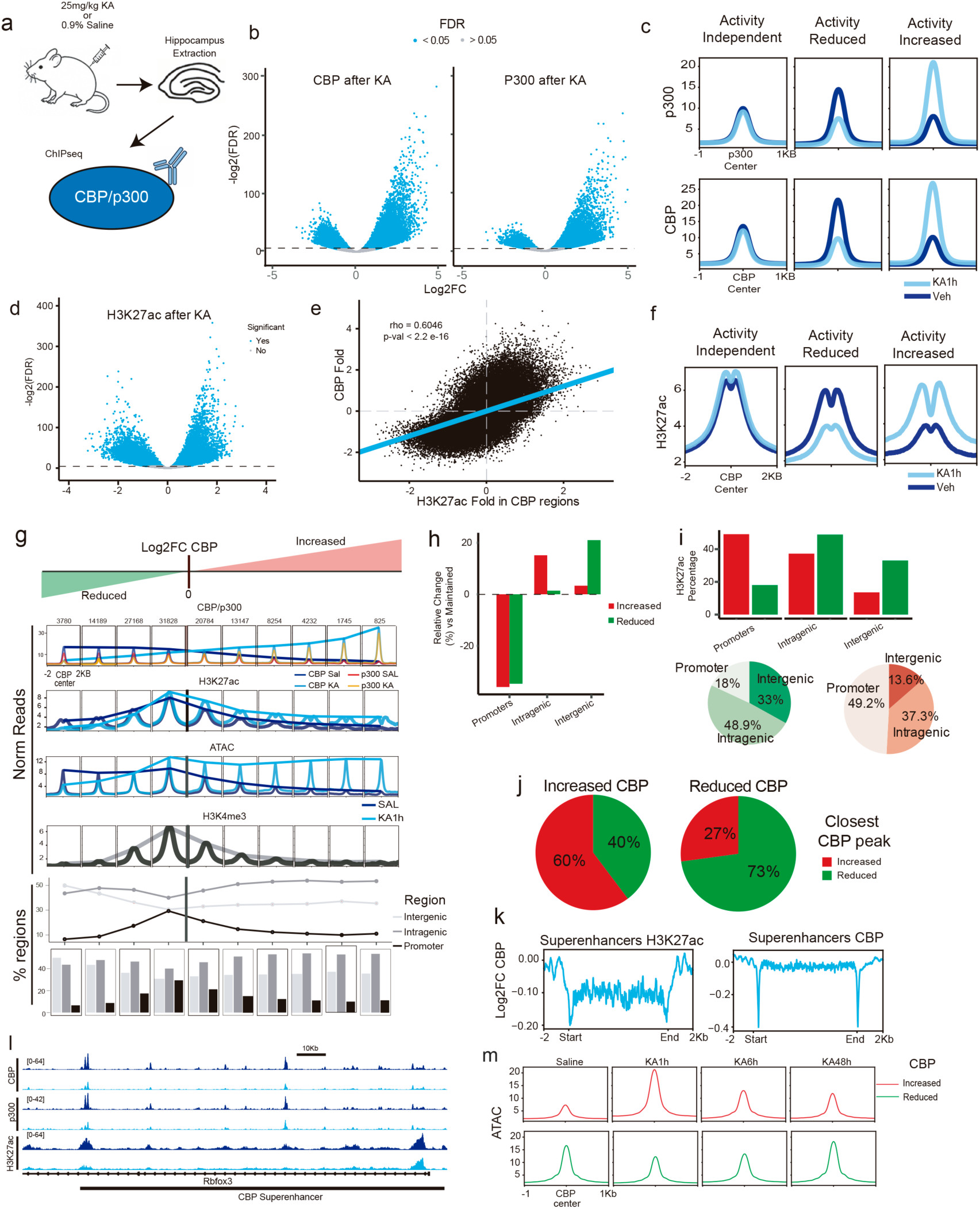
CBP/p300 redistribution alters H3K27 acetylation and chromatin accessibility. **a.** Schematic of the experimental design for multi-omic profiling following KA-induced SE. **b.** Volcano plot showing significant differential binding (DB) peaks for CBP and p300 after KA treatment. **c.** Density plots for three categories of CBP/p300 bound regions: activity-independent, activity-reduced, and activity-increased. **d.** Volcano plot showing differential H3K27ac regions after KA treatment. **e.** Spearman correlation between activity-induced changes in CBP occupancy and H3K27ac signal in CBP-bound regions. **f.** Density plots of H3K27ac signal for the three categories of CBP-bound regions: activity-independent, activity-reduced, and activity-increased. **g.** CBP peaks were split into ten categories according to the magnitude and direction of the KA-induced change. The panels from top to bottom show: the signal used to define the ten categories (CBP and p300 signal in SAL and KA conditions); H3K27ac signal in SAL and KA conditions; ATAC-seq signal in SAL and KA conditions; RNAP2 binding; H3K4me3 signal; H3K4me1 signal; and the percentage of peaks located at promoters, or in intragenic and intergenic regions, shown as line or bar plots. **h.** Relative distribution of CBP-reduced and CBP-increased regions across genomic annotations, normalized to activity-independent regions. **i.** Genomic annotation of regions showing increased or reduced CBP binding or H3K27ac, together with absolute H3K27ac levels in regions with reduced or increased acetylation. **j.** Genomic distances between CBP peaks across categories, showing that peaks tend to lie closer to regions displaying changes in the same direction. **k**. Super-enhancer calling with ROSE using CBP or H3K27ac ChIP-seq data, followed by quantification of activity-dependent changes in CBP occupancy. **l**. Representative example of a super-enhancer showing reduced CBP binding during SE at the *Rbfox3* locus, which encodes the neuronal marker NeuN. **m.** Density plots showing longitudinal ATAC-seq accessibility changes at regions that gain or lose CBP binding.

CBP and p300 behaved very similarly, with substantial overlap in the genomic regions bound by each factor after neuronal activation (**Supp. Fig. S1c-d**), extending our previous findings in resting principal neurons (Lipinski et al., 2020) to the activated state. Overall, ChIP-seq analysis identified over 28,407 regions that lost CBP/p300 binding upon activation, while 20,256 regions exhibited recruitment of these coactivators within 1 h of SE onset (**Fig. 1b-c**). Nevertheless, the majority of CBP/p300-bound regions—over 77,194 regions—remained unaffected by activity.

RNA-seq analyses of bulk hippocampal tissue (**Supp. Fig. S1e-f**) and the cell type-specific transcriptome and translatome of excitatory neurons (Fernandez-Albert et al., 2019) revealed no significant changes in CBP or p300 mRNA levels (**Supp. Fig. S1g**). Likewise, western blotting (**Supp. Fig. S1h-i**) and immunostaining of CA1 (**Supp. Fig. S1j-k**) confirmed that protein levels remained stable after activation, indicating that the observed changes in occupancy reflect redistribution of pre-existing CBP/p300 molecules rather than altered expression.

### KAT3 shuttling dynamically reshapes enhancer landscapes

We next investigated chromatin changes associated with CBP/p300 redistribution. These co-activators are the main acetyltransferases responsible for H3K27ac (Weinert et al., 2018; Lipinski et al., 2020). While several studies have reported increased H3K27ac at activity-regulated loci following neuronal activation in vitro (Malik et al., 2014; Telese et al., 2015), its in vivo regulation remains less well characterized. To address this, we performed H3K27ac ChIP-seq on hippocampal chromatin from control and SE-induced mice (**Supp. Fig. S1l**). We identified approximately 13,800 regions that gained H3K27ac following activation, while 11,600 regions showed a loss of this mark (**Fig. 1d**). Regions gaining CBP/p300 strongly correlated with those acquiring H3K27ac, while regions losing CBP/p300 showed a concordant reduction of H3K27ac (**Figs. 1e-f** and **Supp. Fig. S1m**). These findings suggest a causal relationship between CBP/p300 redistribution and H3K27ac remodeling during neuronal activation.

Given the known association of CBP/p300 and H3K27ac with cis-REs, we next investigated whether their activity-dependent redistribution preferentially affected promoters versus enhancers. To map CBP/p300 redistribution with greater resolution, we divided CBP occupancy changes into 0.5-fold intervals and overlaid signals for several relevant hPTMs (e.g., H3K4me3 and H3K27ac), and neuron-specific chromatin accessibility data (Fernandez-Albert et al., 2019; del Blanco et al., 2024). Regions with little or no change in KAT3 occupancy were marked by strong H3K4me3, consistent with promoters. By contrast, the largest fold changes mapped to H3K4me3-depleted regions with dynamic H3K27ac changes, consistent with enhancers (**Fig.1g**). Overall, CBP/p300- bound regions affected by activity were more likely to be intra- and intergenic enhancers than promoters (**Fig. 1g**, bottom graph). The regions displaying the strongest CBP gains were frequently located within gene bodies (predominantly introns), while regions with the strongest losses mapped into both intragenic and extragenic enhancers (**Fig. 1g**, **Fig. 1h**). Regarding H3K27ac, although activity-driven changes occurred genome-wide, gains were strongly enriched at promoters, whereas losses preferentially occurred at intragenic and intergenic regions (**Fig. 1i**). Interestingly, changes in promoter H3K27ac occurred with comparatively little change in CBP occupancy, whereas H3K27ac gains and losses at enhancers closely tracked corresponding changes in CBP binding (**Supp. Fig. S1n**). These results underscore the enhancer-specific dependence on CBP/p300 for maintaining H3K27 acetylation and suggest that other KATs may contribute to activity-induced H3K27ac gains at promoters.

Notably, CBP losses showed pronounced spatial clustering, with neighboring CBP peaks frequently undergoing concordant reductions (**Fig. 1j**), raising the possibility that these regions correspond to neuronal super-enhancers. To test this possibility, we identified super-enhancers regions using ROSE (Loven et al., 2013; Whyte et al., 2013a) based on H3K27ac or CBP occupancy profiles. Quantification of CBP levels at these super-enhancers confirmed that they undergo CBP depletion following KA-induced activation (**Fig. 1k-l**). Interestingly, we also investigated the set of human neuronal super-enhancers identified by Hnisz and colleagues (Hnisz et al., 2013) and found that after liftOver to the mouse genome, they corresponding regions were also predicted to undergo CBP depletion (**Supp. Fig. S1o**).

Integration with ATAC-seq profiling of chromatin accessibility during and after SE (Fernandez-Albert et al., 2019) revealed substantial overlap between CBP/p300 redistribution and chromatin accessibility changes (**Fig. 1g**). Gains in CBP binding strongly correlated with increases in accessibility, whereas regions losing CBP showed. comparatively modest changes in accessibility. This suggests that regions losing CBP may retain occupancy by basal TFs, while regions gaining accessibility may reflect *de novo* binding of TFs that recruit CBP/p300. Interestingly, increased chromatin accessibility at regions gaining CBP persisted at 6 h and remained detectable 48 h after SE, indicating that a fraction of the activity-induced chromatin response outlasts the initial redistribution of KAT3 proteins (**Fig. 1m**).

Together, these analyses distinguish transcriptional programs that may depend more strongly on CBP/p300 activity from those potentially supported by other acetyltransferase complexes, such as ATAC and SAGA (Nagy and Tora, 2007).

### CBP/p300 shuttling correlates with opposite changes in neuronal plasticity- and identity-related gene programs

We next investigated the relationship between CBP/p300 redistribution and activity-driven transcriptional changes. Gene ontology (GO) enrichment analysis of genes associated with regions showing gains or losses of CBP/p300 binding during SE revealed that losses were associated with genes involved in neuronal differentiation, axonogenesis, and neuronal projection organization, whereas gains were enriched for cytoskeletal remodeling and cellular component organization (**Fig. 2a**).

**Figure 2.**
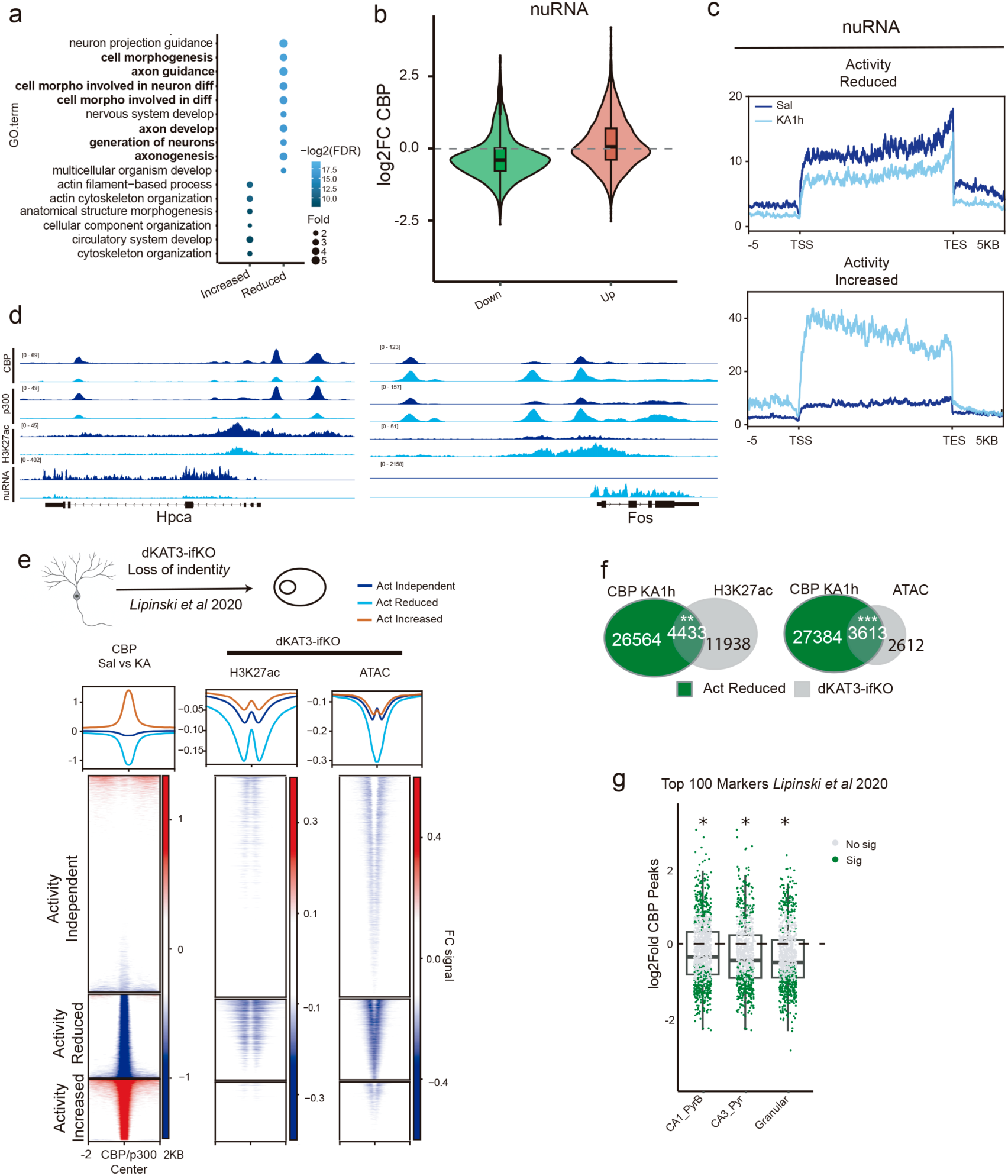
CBP redistribution during neuronal activation correlates with transcriptional changes. **a.** Top 10 Gene Ontology (GO) terms associated with genes linked to regions showing reduced or increased CBP/p300 binding. **b.** CBP occupancy at the top differentially expressed genes (DEGs) identified by nuRNA-seq. **c.** nuRNA metagene profiles for genes associated with regions showing CBP loss or gain and classified as transcriptionally active based on ChromHMM states enriched for H3K27ac and RNAP2. **d.** Genome browser snapshots of representative loci illustrating activity-dependent changes in CBP/p300 occupancy and H3K27ac. **e.** Top: schematic illustrating the loss of neuronal identity features following combined depletion of CBP and p300. Bottom: density plots and heatmaps of CBP-bound regions classified according to the magnitude and direction of CBP changes in the KA model. The corresponding H3K27ac and chromatin accessibility (ATAC-seq) changes at these regions in dKAT3-ifKO neurons are shown alongside. **f.** Venn diagrams showing the overlap between regions losing CBP in the KA model and regions showing loss of H3K27ac or chromatin accessibility in the dKAT3-ifKO model. **g.** CBP Log2 fold changes at CBP peaks associated with the top 100 marker genes for hippocampal excitatory neuron populations defined by snRNAseq (Lipinski et al., 2020), comparing SAL and KA conditions.

To better understand how CBP binding relates to transcriptional changes, we compared activity-driven changes in CBP occupancy with our analyses of activity-dependent changes in nuclear RNA (nuRNA-seq) (Fernandez-Albert et al., 2019), mature cytoplasmic mRNA (this study), and ribosome-associated RNA (Fernandez-Albert et al., 2019). Across all datasets, activity-driven changes in CBP occupancy correlated with changes in transcript abundance, with the association being strongest for nuclear RNA and progressively weaker for mature and ribosome-associated RNA (**Supp. Fig. S2a**). Moreover, decreased CBP binding was associated with gene downregulation only in the nuRNA-seq dataset (**Supp. Fig. S2b**). Consistently, differentially expressed genes (DEGs) identified by nuRNA-seq analysis exhibited CBP gain at upregulated genes and CBP loss at downregulated genes (**Fig. 2b**). Analysis of transcribed regions defined by ChromHMM based on enrichment for CBP, H3K27ac, and RNAP2 (**Supp. Fig. S2c**) confirmed reduced transcription at regions losing CBP and increased transcription at regions gaining CBP (**Fig. 2c**). This pattern likely reflects the rapid detection of activity-dependent transcriptional changes in nuclear RNA, whereas reductions in mature mRNA abundance or ribosome-associated transcripts are delayed by transcript degradation kinetics.

Analysis based on CBP occupancy at transcription start sites (TSSs) or associated enhancers further revealed a strong correlation between activity-driven CBP recruitment and gene induction (**Supp. Fig. S2d**). The promoters of representative activity-induced genes gained both CBP/p300 binding and H3K27ac, including IEGs such as *Fos* and *Arc*, and late-response genes such as *Bdnf, Scg2* and *Nptx2* (**Fig. 2d** and **Supp. Fig. S2e-f**). Conversely, promoters of activity-downregulated genes lost both CBP/p300 binding and H3K27ac. Among these genes were numerous neuronal markers (e.g., *Slc17a7*, encoding VGLUT1), and genes with functions in synapse formation and dendritic arborization (e.g., *Foxo6, Wnt7b*) (**Fig. 2d** and **Supp. Fig. S2e-f**). CBP redistribution was also evident at enhancers associated with differentially expressed genes (**Supp. Fig. S2g**). Representative loci further showed concordant changes in enhancer transcription and CBP/p300 occupancy (**Supp. Fig. S2h**).

We have previously shown that CBP and p300 are jointly required to maintain neuronal identity, and that their combined ablation in adult neurons causes a rapid loss of neuronal features accompanied by H3K27 deacetylation (Lipinski et al., 2020). Other studies have suggested a broader role for CBP/p300 in identity maintenance in photoreceptors, muscle, kidney, and pancreatic endocrine cells (Pentz et al., 2012; Hennig et al., 2013; Fang et al., 2014; Svensson et al., 2020; Zhang et al., 2021a; Wang et al., 2026). Based on these findings and the pattern described above, we hypothesized that neuronal activity promotes the redistribution of CBP/p300 from genes supporting neuronal identity—typically large genes located within or near super-enhancers—to activity-induced genes, which tend to include shorter, rapidly transcribed genes and are frequently associated with distal activity-regulated enhancers (Saha et al., 2011; Malik et al., 2014). Consistent with this hypothesis, regions that lose CBP following SE were particularly sensitive to genetic ablation of CBP/p300, exhibiting pronounced H3K27 deacetylation and reduced chromatin accessibility in double conditional knockout neurons (dKAT3-ifKOs; Lipinski et al., 2020) (**Fig. 2e**). Moreover, regions losing CBP during SE significantly overlapped those showing H3K27ac or accessibility loss following KAT3 ablation (**Fig. 2f**). These observations suggest that regulatory regions supporting neuronal identity are particularly dependent on KAT3 activity and transiently lose KAT3 occupancy during neuronal activation. To further test this idea, we examined the association between changes in CBP binding and the top 100 excitatory neuron marker genes defined by snRNA-seq studies in the hippocampus (Lipinski et al., 2020). These markers showed marked CBP depletion following KA treatment (**Fig. 2g**). Together, these findings support a model in which CBP/p300 dynamically redistribute from KAT3-dependent neuronal identity-associated regulatory regions toward activity-induced loci, accompanied by reciprocal changes in H3K27ac, chromatin accessibility, and transcriptional output.

### KAT3 proteins shuttle between NeuroD and AP-1 binding sites

Since neither CBP nor p300 bind directly to DNA (Kwok et al., 1994; Lundblad et al., 1995), we next asked which specific TFs could be involved in their redistribution after neuronal activation. Motif analysis using HOMER revealed strong enrichment for AP-1 and CREB binding motifs upon neuronal activation (TPA response elements, TREs, and cAMP response element, CREs, respectively) at regions that gained CBP binding, whereas E-box motifs, including a NeuroD-family motif, were enriched at regions that lost CBP (**Fig. 3a** and **Supp. Fig. S3a**). Since NeuroD family members (e.g., NeuroD2 and NeuroD6) act as terminal selectors, maintaining neuronal identity programs throughout life (Deneris and Hobert, 2014), their enrichment at cis-REs losing CBP/p300 during SE further supports the model of transient attenuation of neuronal identity-associated regulatory programs during neuronal activation. Interestingly, many regions with stable CBP/p300 binding harbored both E-box and TRE motifs, suggesting that motif coexistence may buffer them from activity-driven redistribution. Consistent with this possibility, TRE and CRE motifs were modestly depleted from CBP-lost regions relative to background (**Supp. Fig. S3a**). Notably, motif enrichment at differentially acetylated regions mirrored these results: regions losing H3K27ac were enriched for E-box motifs, whereas regions gaining H3K27ac were enriched for CRE and TRE motifs (**Supp. Fig. S3b**).

**Figure 3.**
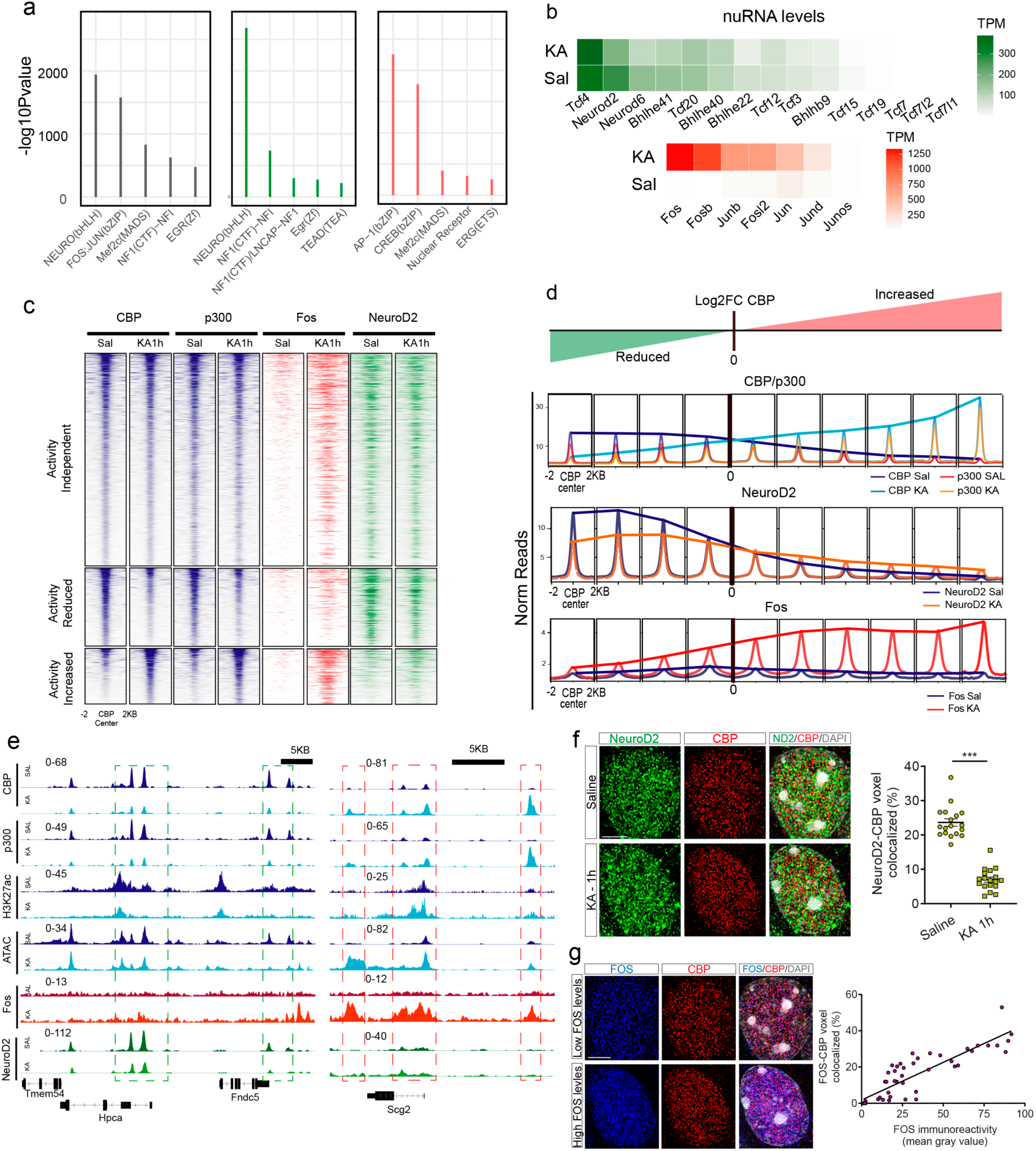
Activity-induced AP-1 recruitment redistributes CBP/p300 away from NeuroD-associated loci. **a.** Motif analysis across the three CBP region categories. bHLH E-box motifs were enriched in activity-reduced and activity-independent regions, whereas AP-1 TRE motifs were enriched in activity-increased and activity-independent regions. CRE motifs is also enriched in activity-increased regions. **b.** Expression of NeuroD- and AP-1-family transcription factors in principal neurons of the adult mouse hippocampus under saline and 1 h post-KA conditions. **c.** Heatmaps of hippocampal ChIP-seq signal under saline and 1 h post-KA conditions for CBP, p300, FOS, and NeuroD2. **d.** CBP-bound regions were divided into 0.5-fold change intervals, and corresponding CBP, p300, FOS, and NeuroD2 ChIP-seq profiles were plotted across the resulting categories. **e.** Genome browser snapshots of representative enhancer loci illustrating CBP/p300 redistribution together with FOS and NeuroD2 occupancy and associated chromatin changes. **f.** Representative super-resolution and deconvolved images illustrating spatial colocalization between CBP and NeuroD2 within the nuclei of CA1 hippocampal neurons under saline conditions and 1 h after KA administration. Right: quantification of CBP–NeuroD2 voxel co-labeling, showing reduced spatial overlap following KA treatment. n= 18 cells/3 mice per condition. Scale bar: 5 μm. **g.** Representative super-resolution and deconvolved images illustrating spatial colocalization between CBP and FOS within CA1 neuronal nuclei following neuronal activation. Right: positive correlation between CBP–FOS voxel co-labeling and nuclear FOS levels. N = 50 cells/12 mice. Scale bar: 5 μm.

To further explore activity-dependent motif occupancy, we performed digital footprinting with ATAC-seq. Regions gaining CBP showed marked increases in AP-1 (FOS:JUN) footprinting after KA, whereas NeuroD-family footprints changed only modestly at regions losing CBP (**Supp. Fig. S3c**). This suggests that NeuroD family members largely remain associated with chromatin during stimulation, whereas AP-1 family factors are newly recruited to activity-induced regions.

Together, these results support a model in which bHLH and AP-1/CREB TFs differentially engage a limited pool of CBP/p300 during neuronal activation. Mechanistically, this is consistent with the modular structure of CBP/p300, which contains multiple interaction domains that mediate contacts with distinct TF families, including CREB, AP-1 factors, and bHLH proteins (Goodman and Smolik, 2000; Lipinski et al., 2019). We therefore hypothesized that neuronal activation shifts CBP/p300 occupancy away from NeuroD-associated loci toward newly engaged AP-1/CREB-regulated regions, thereby redistributing co-activator availability from neuronal identity programs to activity-dependent plasticity genes.

To empirically test our *in silico* predictions, we assessed the occupancy of hippocampal chromatin by two prominent members of the AP-1 and NeuroD families under basal conditions and 1 h after KA treatment using ChIP-seq. For AP-1, we focused on FOS, given its robust induction by KA and widespread use as a marker of neuronal activation (**Fig. 3b**). For NeuroD TFs, we focused on NeuroD2 because of its critical role as a terminal selector in excitatory neurons (Deneris and Hobert, 2014; Tutukova et al., 2021), and its strong expression in hippocampal principal neurons (**Fig. 3b**). ChIP-seq analysis using specific antibodies against each protein revealed striking differences in their behavior during SE. FOS displayed dramatic activity-induced binding across the genome, while NeuroD2 binding remained largely stable (**Fig. 3c**). These results suggest that CBP/p300 redistribution is not primarily driven by loss of NeuroD2 occupancy from chromatin, but instead coincides with strong activity-induced FOS binding.

To link TF occupancy with CBP dynamics, we stratified CBP peaks according to the strength of gain or loss after KA as shown in **Fig. 1g**. The strongest CBP losses were associated with the highest NeuroD2 occupancy, whereas increasing CBP recruitment tracked progressively stronger FOS binding (**Fig. 3d**). Analysis of combinatorial TF occupancy further showed that FOS/NeuroD2 co-bound regions underwent substantially smaller CBP changes than regions bound by either factor alone (**Supp. Fig. S3d-e**). At baseline, NeuroD2 showed extensive co-occupancy with CBP. After KA, CBP occupancy at NeuroD2-bound sites decreased—unless FOS was also present, in which case CBP loss was strongly attenuated, indicating that simultaneous NeuroD2 and FOS occupancy largely neutralized the directional change in CBP binding observed at sites occupied predominantly by either TF. Conversely, FOS binding in the absence of NeuroD2 was associated with robust CBP recruitment. Representative loci illustrating these dynamics are shown in **Fig. 3e**.

Immunostaining analyses confirmed that nuclear levels of CBP and NeuroD2 remained unchanged, whereas FOS was strongly induced by KA (**Supp. Fig. S3f**). To directly visualize these events at the single-neuron level, we used super-resolution microscopy to explore the spatial distribution of these proteins within the nuclei of CA1 pyramidal neurons. We quantified their overlap by measuring the proportion of voxels from one protein that contained signal from the other. In line with the ChIP-seq results, CBP colocalized extensively with NeuroD2 at baseline, but this spatial association sharply decreased 1 h after KA (**Fig. 3f**). In contrast, CBP colocalization with FOS increased upon neuronal activation and scaled with nuclear FOS abundance (**Fig. 3g**). Time-course analysis revealed reciprocal and transient changes in CBP association with NeuroD2 and FOS: NeuroD2–CBP colocalization decreased as FOS–CBP colocalization increased, with both returning toward basal levels at later time points (**Supp. Fig. S3g**).

Together, these results show that NeuroD2 remains largely associated with chromatin while CBP occupancy at NeuroD2-bound regions decreases during neuronal activation, whereas strong activity-induced FOS binding is accompanied by robust CBP recruitment. The reciprocal temporal changes in NeuroD2–CBP and FOS–CBP colocalization further support a model in which neuronal activation redistributes KAT3 coactivators between neuronal identity- and activity-associated transcriptional networks. Thus, AP-1 factors, and in particular FOS, emerge as major candidates driving the activity-dependent redistribution of CBP/p300.

### CBP/p300 redistribution remodels activity-dependent chromatin interactions

CBP and p300 have been implicated in genome organization and DNA conformation (Wong et al., 2014), but their specific contribution to 3D chromatin organization remains poorly understood. We reasoned that activity-dependent changes in CBP/p300 association with TFs and chromatin could affect enhancer–promoter communication and higher-order genome organization. To address this possibility, we integrated CBP ChIP-seq data with our previously generated Hi-C datasets from glutamatergic hippocampal neurons under saline and KA conditions (Fernandez-Albert et al., 2019). Aggregate Hi-C maps revealed that CTCF peaks coincided with sharply structured chromatin boundaries, whereas CBP peaks were associated with broader local contact enrichment rather than canonical insulating architecture (**Supp. Fig. S4a**).

We next classified CBP-associated regions into three categories according to their behavior during SE, as described in **Fig. 1**: maintained, lost, or gained. Aggregate Hi-C maps from saline and KA conditions, centered on these regions, revealed a cross-shaped pattern of local contact enrichment in all three groups, indicating that regions associated with dynamic CBP binding are embedded in interaction-rich chromatin environments already under basal conditions and that their overall local contact architecture is not grossly altered during SE. As expected, maintained and lost sites, both of which are occupied by CBP in saline-treated animals, displayed this pattern. Remarkably, regions that gained CBP following KA stimulation also exhibited broad local contact enrichment at baseline, suggesting that activity-dependent CBP recruitment preferentially occurs at loci with a pre-existing interaction-rich 3D chromatin environment (**Supp. Fig. S4b)**.

To examine enhancer–promoter interactions at higher resolution, we performed H3K27ac HiChIP on hippocampi from saline- or KA-treated mice (**Fig. 4a-b** and **Supp. Fig. S4c**). H3K27ac HiChIP enriches for contacts involving active chromatin, enabling sensitive detection of short-range regulatory interactions while recapitulating the major features of overall 3D genome organization (**Supp. Fig. S4d**). FitHiChIP analysis identified approximately 120,000 loops anchored at H3K27ac-enriched regions. Differential analysis revealed ∼10,000 loops whose interaction strength changed following KA treatment, with both gained and lost contacts (**Fig. 4c**). Most H3K27ac-mediated contacts overlapped with CBP binding, and interaction strength increased with CBP occupancy (**Supp. Fig. S4e)**. Moreover, regions involved in contacts that weakened after KA tended to lose CBP, whereas regions involved in strengthened contacts tended to gain CBP (**Fig. 4d**), linking CBP redistribution to activity-dependent remodeling of chromatin interactions.

**Figure 4.**
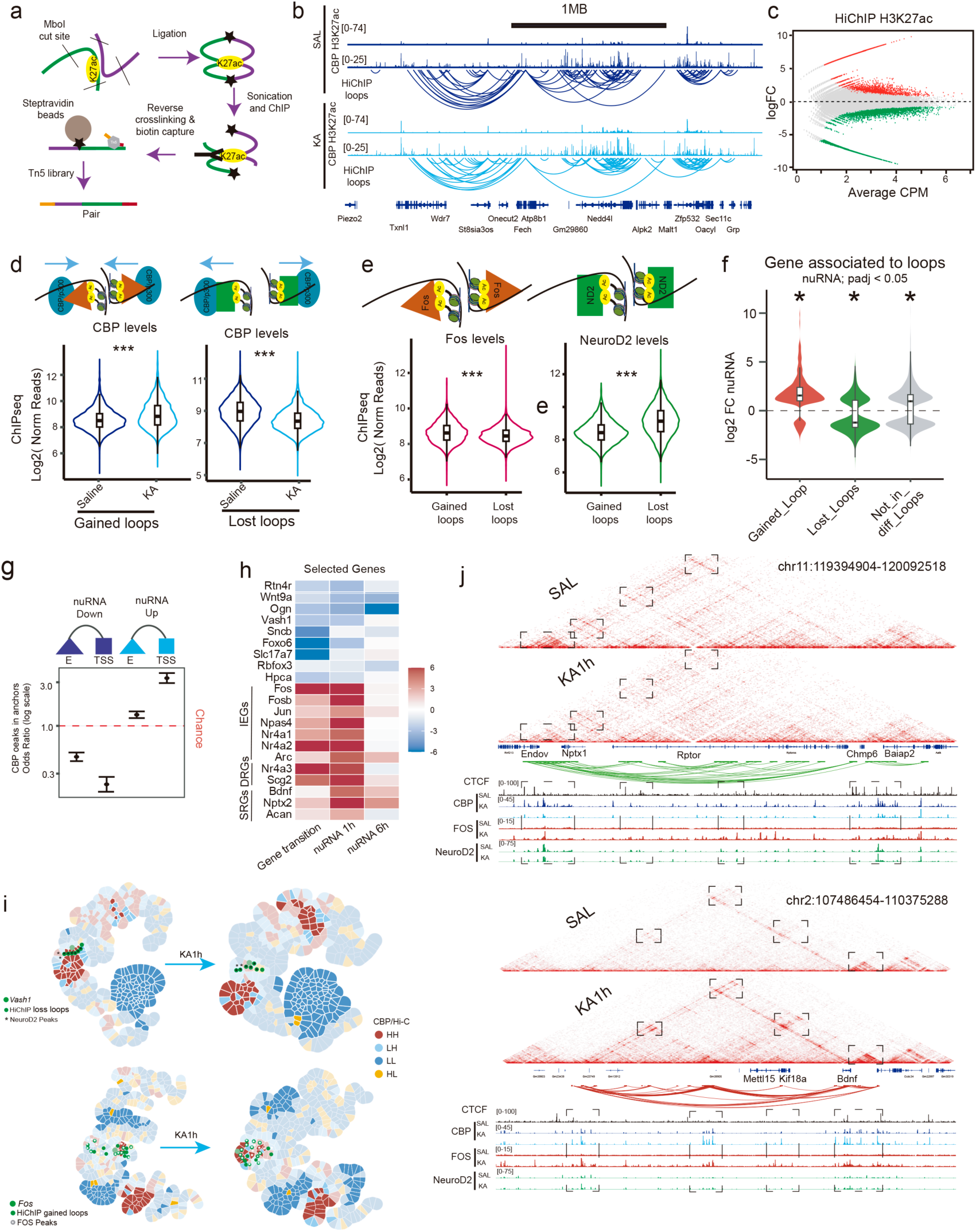
Neuronal activation remodels CBP-associated chromatin loops in a TF-dependent manner. **a.** Schematic of H3K27ac HiChIP experiment. **b.** Representative genomic region illustrating H3K27ac HiChIP loops and their overlap with CBP and H3K27ac peaks. **c.** MA plot showing differential HiChIP loops following KA treatment. **d.** Aggregate HiChIP signal at differential loops intersecting regions with activity-induced changes in CBP binding. **e.** Left: FOS ChIP-seq signal after KA treatment at differential loops. Right: NeuroD2 ChIP-seq signal under basal conditions at differential loops. **f.** nuRNA expression changes for DEGs associated or not associated with differential loops. nuRNA data from (Fernandez-Albert et al., 2019). **g.** Enrichment of CBP gain or loss at TSSs and putative enhancers associated with DEGs, irrespective of detectable loop remodeling. Enhancer anchors were defined as HiChIP anchors not overlapping a TSS. E = putative enhancer. nuRNA data from (Fernandez-Albert et al., 2019). **h.** Examples of spatial-state transitions for neuronal identity and activity-regulated genes during SE. **i.** CBP signal mapped onto 3D chromatin graphs and represented as Gaudí plots, with each genomic bin classified according to its local Moran’s I (LMI) category; solid colors indicate statistically significant assignments (P < 0.05). Representative activity-induced remodeling of HH metaloci at *Vash1* and *Fos* is shown. HiChIP differential loops and peaks of the ChIPseq from NeuroD2 and FOS near these genes are marked. **j.** Juicer and IGV views of representative loci showing loss (top) or gain (bottom) of chromatin interactions, together with HiChIP, CTCF, CBP, FOS, and NeuroD2 profiles. CTCF ChIP-seq data are from (Shen et al., 2012); nuRNA data are from (Fernandez-Albert et al., 2019); FOS ChIP-seq are from (Alaiz-Noya et al., 2025).

We next asked whether these activity-regulated contacts were characteristic of differentiated neurons. Using H3K27ac profiles from neural progenitor cells (NPCs) and cortical neurons (CNs) (Bonev et al., 2017), we found that most regions participating in activity-dependent loops were preferentially marked in CNs, indicating that these regulatory elements are largely neuron specific (**Supp. Fig. S4f**). This pattern is consistent with the expression of transcription factors involved in neuronal identity and maturation, including proneural NEUROD factors, which are more highly expressed in CNs than in NPCs (Bonev et al., 2017) (**Supp. Fig. S4g**).

Because differential HiChIP loops can arise from changes in H3K27ac abundance or from altered contact frequency independently of local acetylation, we next dissected these two contributions. Approximately 40% of differential loops were not accompanied by significant changes in H3K27ac, indicating that a substantial fraction of activity-dependent interaction remodeling cannot be explained solely by changes in local acetylation (**Supp. Fig. S4h**). Together with the corresponding changes in CBP occupancy, these results suggest that TF-associated CBP redistribution may contribute to enhancer–promoter rewiring even in the absence of detectable local H3K27ac changes. Consistent with this hypothesis, aggregate HiChIP analysis confirmed the expected changes in interaction signal at loci involved in differential H3K27ac HiChIP loops that also intersected regions showing differential CBP binding (**Supp. Fig. S4i**). Importantly, most of these interactions were also detectable in the corresponding Hi-C datasets, although with weaker signal because of the lower sensitivity of Hi-C for active regulatory contacts (**Supp. Fig. S4j**). These results support the conclusion that CBP redistribution and H3K27ac-associated regulatory activity accompany remodeling of intra-TAD interactions during neuronal activation.

To assess the contribution of TFs to these architectural changes, we examined the overlap between differential loops and NeuroD2 and FOS binding. Regions involved in interactions that weakened following KA were enriched for NeuroD2, whereas regions participating in newly gained or strengthened interactions were strongly associated with FOS (**Fig. 4e**). Given that bHLH TFs participate in chromatin contacts associated with neuronal maturation (Bonev et al., 2017; Winick-Ng et al., 2021), whereas AP-1 factors promote activity-dependent chromatin interactions (Fernandez-Albert et al., 2019; Beagan et al., 2020), these results further support a shift between 3D regulatory configurations associated with neuronal identity and activity-dependent transcriptional programs.

We sought independent support for this model using Genome Architecture Mapping (GAM), a ligation-free approach that infers chromatin contacts from the co-segregation frequencies of genomic regions in nuclear slices. GAM avoids biases associated with proximity ligation and is particularly sensitive to interactions among active regions and to multi-way contacts, thereby enabling the detection of potential regulatory hubs (Beagrie et al., 2017). We examined the top 10% of strongest CA1 pyramidal neuron-specific interactions identified by GAM at 20 kb resolution (Winick-Ng et al., 2021). These interactions confirmed enrichment of NeuroD2 binding at baseline CBP-associated regions (**Supp. Fig. S4k**), independently supporting our HiChIP results and extending them to higher-order interactions.

### CBP/p300-dependent 3D remodeling is coupled to transcriptional regulation

Several studies have linked changes in chromatin conformation to transcriptional regulation (Fernandez-Albert et al., 2019; Beagan et al., 2020; Stroud et al., 2020; Calderon et al., 2022). We therefore asked whether activity-dependent loop remodeling was associated with the bidirectional transcriptional changes observed during SE. For most genes associated with differential loops, the direction of expression change matched the direction of interaction remodeling, supporting a relationship between transcriptional regulation and chromatin reorganization (**Fig. 4f**). However, a subset of transcriptional changes occurred without detectable loop remodeling and was nevertheless associated with altered CBP occupancy. Downregulated genes were enriched for CBP loss at both TSSs and associated enhancers, whereas upregulated genes showed the strongest enrichment for CBP gain at TSSs (**Fig. 4g**). Consistently with the pre-existing local contact enrichment observed at CBP-gain regions (**Supp. Fig. S4b**), the comparatively modest increase in CBP occupancy at enhancers suggests that some activity-regulated genes may engage enhancer configurations that are already established before stimulation. Other enhancer–promoter interactions, by contrast, emerge or become strengthened specifically in response to stimulation.

To characterize these changes at the level of 3D regulatory neighborhoods, we applied METALoci, a framework that uses spatial autocorrelation to determine how the distribution of a genomic feature—in this case CBP occupancy—is related to 3D chromatin organization at both local and global scales (Mota-Gómez et al., 2026). METALoci classifies each genomic bin according to both its own signal and that of its spatially proximal neighbors. High–high (HH) bins contain high CBP signal and are surrounded in 3D space by other CBP-enriched bins, whereas low–low (LL) bins show low CBP signal both locally and in their spatial neighborhood. High–low (HL) bins contain high CBP signal within a low-signal neighborhood, whereas low–high (LH) bins contain low CBP signal within a high-signal neighborhood. Contiguous groups of statistically significant HH bins are defined as *metaloci* and represent spatially organized regions enriched for the feature under investigation, which in our analysis corresponds to CBP occupancy.

For each gene in the mouse genome, we used METALoci to reconstruct these 3D regulatory neighborhoods under basal and SE conditions by integrating Hi-C contacts with CBP ChIP-seq signal. In this context, HH metaloci can be interpreted as CBP-enriched 3D regulatory hubs. Although the overall numbers of genes assigned to HH and LL categories changed only modestly during SE, a substantial number of individual genes switched spatial states. Genes transitioning from HL or LL states toward HH exhibited coordinated transcriptional changes (**Fig. 4h**). Intriguingly, among the top DEGs identified 6 h after KA treatment, these spatial transitions were already detectable at 1 h, suggesting that reorganization of 3D regulatory neighborhoods may precede the full transcriptional response of the corresponding loci (**Supp. Fig. S4l-m**). Thus, METALoci revealed extensive rewiring of CBP-enriched regulatory hubs during SE, affecting the 3D chromatin environments of hundreds of genes. Representative METALoci maps illustrate remodeling of CBP-enriched 3D regulatory neighborhoods at *Vash1* and *Fos* (**Fig. 4i**).

Additional examples illustrate the functional relevance of this remodeling. *Nptx1*, a gene implicated in memory and glutamatergic transmission (Cummings et al., 2017; Jin et al., 2025) showed loss of chromatin interactions following neuronal activation (**Fig. 4j, upper panel**). Conversely, *Bdnf*, a late activity-regulated gene encoding a neurotrophin with a critical role in neuronal plasticity (Patterson et al., 1996), gained regulatory interactions (**Fig. 4j, lower panel**).

Together, these findings show that activity-dependent redistribution of CBP/p300 is closely associated with remodeling of intra-TAD regulatory interactions. CBP/p300 redistribution therefore extends beyond changes in enhancer acetylation and accompanies the reorganization of long-range enhancer–promoter contacts. These results link TF-dependent co-activator redistribution to dynamic changes in 3D chromatin organization and suggest a structural mechanism through which neuronal state transitions can be coordinated with changes in gene expression.

### KAT3 shuttling is also observed in response to physiological stimuli

To determine whether the changes observed during SE also occur during physiological activation of hippocampal principal neurons, we performed CBP ChIP-seq in the hippocampus of animals exposed to a novel environment (NE) (**Fig. 5a**). Bulk hippocampal CBP ChIP-seq after NE exposure showed the same directional redistribution observed during SE, but with substantially smaller effect sizes. This difference likely reflects the reduced number of activated neurons and the lower intensity of activation following physiological stimulation compared with SE (**Fig. 5b**). Consistent with these results, reanalysis of ATAC-seq data in FOS⁺ and FOS⁻ neurons after NE (Fernandez-Albert et al., 2019) concordant accessibility changes at SE-defined CBP-reduced and CBP-increased regions following NE (**Fig. 5c**).

**Figure 5.**
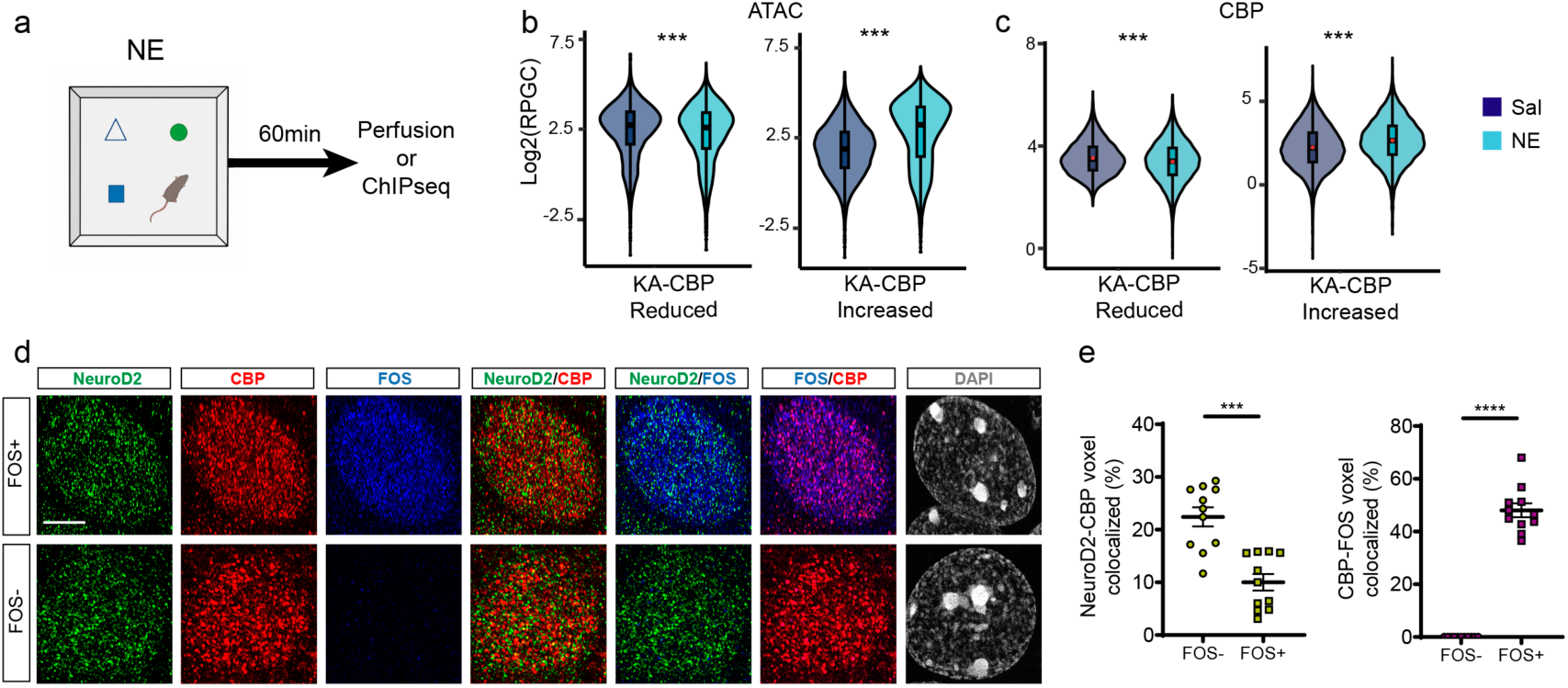
KAT3 shuttling is also observed during physiological neuronal activation. **a.** Schematic of the experimental design showing CBP ChIP-seq or tissue collection for imaging 1 h after novel environment (NE) exposure. **b.** CBP signal following NE exposure at regions classified according to CBP redistribution during SE. **c.** ATAC-seq signal in FOS⁺ and FOS⁻ neurons following NE exposure at regions classified according to CBP redistribution during SE. **d.** Representative super-resolution and deconvolved images showing CBP colocalization with NeuroD2 and FOS within the nuclei of CA1 hippocampal neurons following NE exposure, comparing FOS⁻ and FOS⁺ neurons from the same animals. **e.** Quantification of voxel co-labeling showing reduced CBP–NeuroD2 colocalization (left) and increased CBP–FOS colocalization (right) in FOS⁺ compared with FOS⁻ neurons following NE exposure. n = 11 cells/4 mice per condition. Scale bar: 5 μm.

To further confirm that CBP redistribution during physiological activation mirrors that observed after KA, we examined CBP colocalization with NeuroD2 and FOS in the nuclei of FOS⁺ and FOS⁻ CA1 pyramidal neurons from mice exposed to NE using super-resolution microscopy. The analysis revealed that activity-induced FOS strongly colocalized with CBP, while CBP–NeuroD2 colocalization was reduced in FOS⁺ neurons compared to FOS⁻ neurons (**Fig. 5d-e**), supporting the occurrence of KAT3 shuttling during physiological neuronal activation.

### KAT3 shuttling also occurs in cultured neurons upon depolarization

To further dissect the temporal dynamics and molecular mechanisms of KAT3 shuttling during neuronal activation, we turned to primary hippocampal neuronal cultures (PNCs). Activity-dependent transcription induced by KCl treatment in PNCs (**Fig. 6a**) recapitulated key features of the in vivo response, including the concomitant upregulation of early and late response genes (**Fig. 6b-c**) and the downregulation of neuronal identity genes (**Fig. 6d**). Importantly, repression was considerably more pronounced when measured using intronic transcripts than using exon–exon amplicons, indicating that the immediate effect is most readily detected at the level of ongoing transcription, consistent with the transient nature of the effect observed *in vivo*. Notably, the impact on mature mRNAs became broader and more evident 3 h after KCl treatment (**Fig. 6e**). **Fig. 6f** shows some representative examples. Furthermore, with prolonged stimulation, the transcriptional response eventually propagated to the protein level, as reflected by reduced NEUROD2 immunoreactivity after 24 h of KCl treatment (**Fig. 6g-h**), suggesting that sustained activation can extend the transcriptional consequences of KAT3 redistribution to more persistent changes in neuronal gene expression.

**Figure 6.**
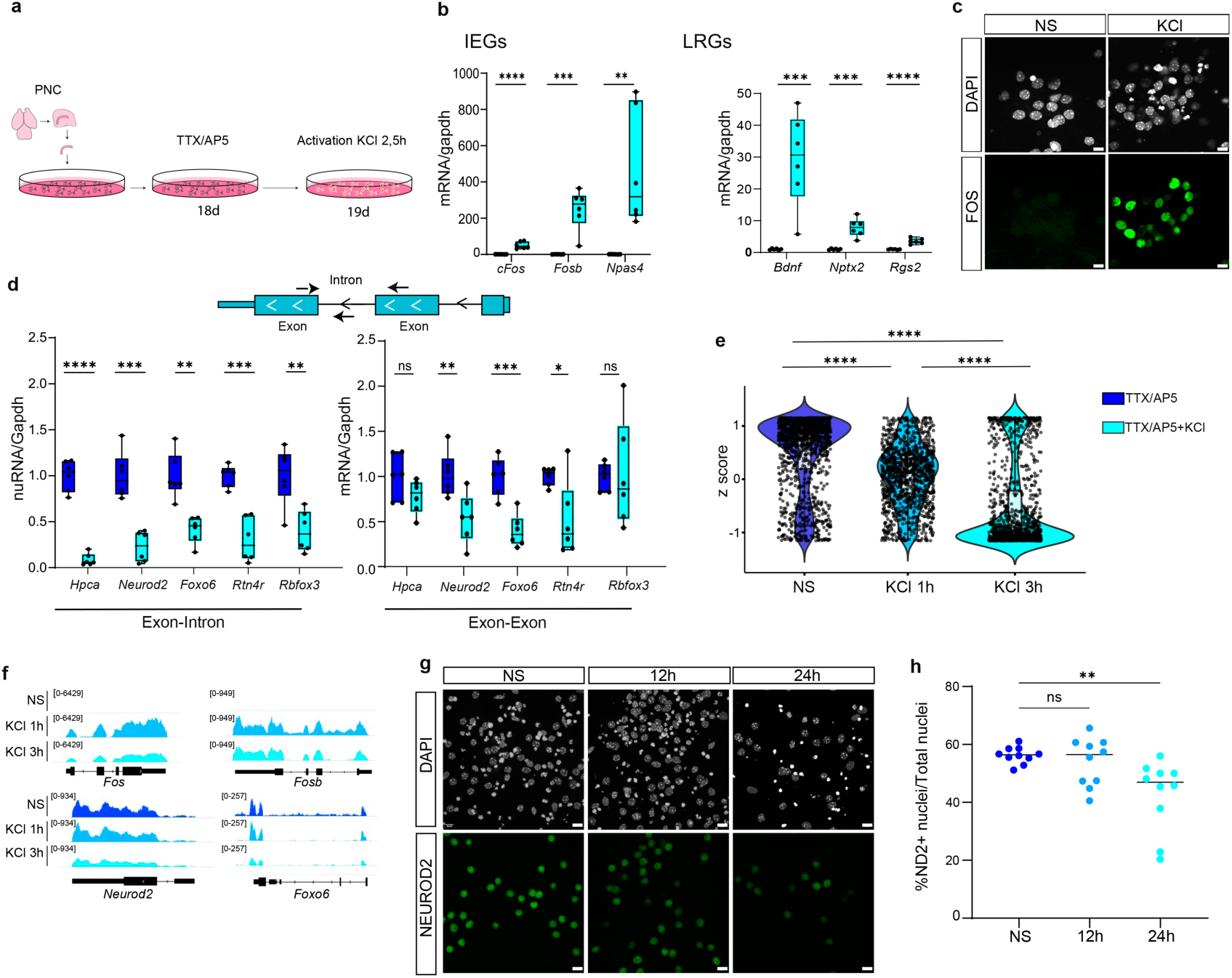
Sustained neuronal activation leads to downregulation of identity markers. **a.** Experimental design for activity induction in primary hippocampal neurons. Primary hippocampal neurons derived from E17 tissue were cultured to DIV14 and pharmacologically silenced before KCl stimulation. **b.** Representative image showing robust induction of FOS following stimulation. Scale bar: 10 μm. **c.** RT–qPCR analysis showing robust induction of early- and late-response gene programs, with first- and second-wave activity-regulated genes significantly upregulated (n = 6; mean ± SEM; two-tailed t-test: ****p < 0.0001, ***p < 0.001, **p < 0.01). **d.** Activity-dependent repression of neuronal identity programs. Top: schematic overview of the primer design used to distinguish changes in ongoing transcription from mature transcript abundance. Bottom: RT–qPCR showing reduced expression of neuronal identity genes following stimulation, with intronic and exon–exon transcript measurements analyzed separately (n = 6; mean ± SEM; two-tailed t-test: ****p < 0.0001, ***p < 0.001, **p < 0.01, *p < 0.05). **e.** RNA-seq log₂ fold-change distribution showing progressively broader downregulation of neuronal identity genes following prolonged activation (Kruskal–Wallis test, p < 2.2 × 10⁻⁶). **f.** RNA-seq tracks illustrating induction of CBP-gain genes (Fos, Fosb) and repression of CBP-loss targets (Neurod2, Foxo6). **g.** Immunostaining time course of NeuroD2 at 0, 12, and 24 h, showing a gradual reduction in signal intensity during sustained activation. Scale bar: 10 μm. h. Quantification of NeuroD2 immunoreactivity over time, confirming significant reduction following prolonged stimulation (one-way ANOVA).

### FOS expression is sufficient to mimic the activity-dependent repression of neuronal identity genes

The observation that KAT3 shuttling also occurs in PNCs provided an experimental system in which to dissect the underlying mechanism using loss-of-function (LOF) and gain-of-function (GOF) approaches.

Efficient repression of NeuroD2 using CRISPR-KRAB (**Fig. 7a-b** and **Supp. Fig. S5a**) did not prevent the downregulation of neuronal identity genes (**Fig. 7c**) or alter IEG induction (**Supp. Fig. S5b**). Thus, NeuroD2 reduction alone is not sufficient to block activity-dependent repression of identity genes. Heterologous NeuroD2 overexpression (**Fig. 7d-e** and **Supp. Fig. S5c**) did not rescue identity gene downregulation either (**Fig. 7f**) and did not interfere with IEG induction (**Supp. Fig. S5d**), indicating that elevated NeuroD2 levels are insufficient to preserve the identity-associated transcriptional program during stimulation. These results indicate that NeuroD2 abundance is neither necessary nor sufficient to determine the activity-dependent repression of these neuronal identity genes. This may reflect functional redundancy among NeuroD or other bHLH family members.

**Figure 7.**
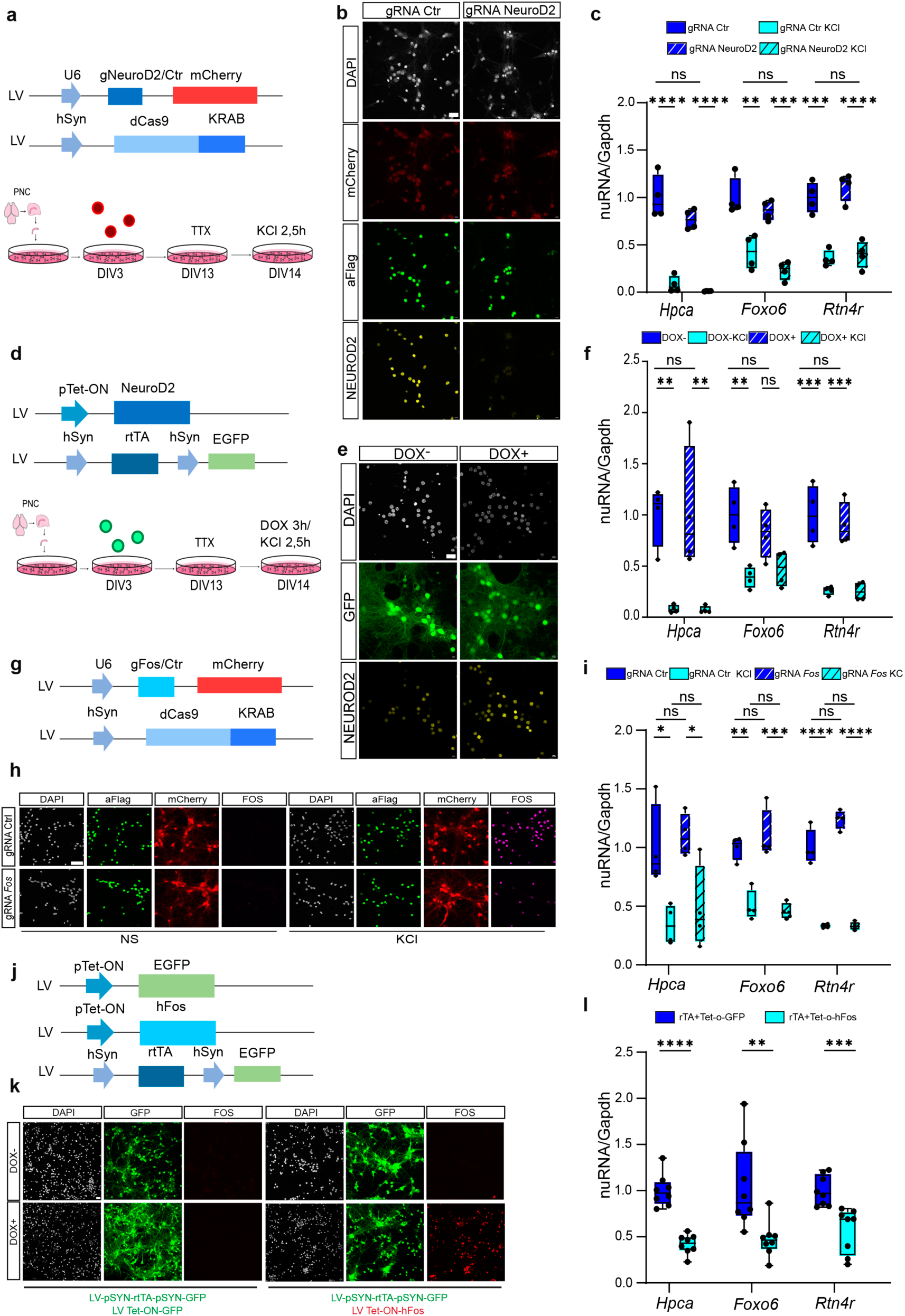
FOS expression is sufficient to mimic activity-dependent repression of neuronal identity genes. **a-c.** NeuroD2 loss-of-function analysis. **a.** Lentiviral delivery of the dCas9-KRAB–MeCP2 system with NeuroD2-targeting guide RNAs was used to repress NeuroD2 in PNCs infected at DIV3; cultures were maintained until DIV14 before KCl-induced activation. **b.** Protein-level validation confirms efficient NeuroD2 repression. Scale bar: 10 μm. **c.** RT–qPCR analysis of activity-repressed identity genes showing that NeuroD2 depletion does not prevent their activity-dependent downregulation (n = 4; mean ± SEM; one-way ANOVA). **d-f.** NeuroD2 gain-of-function analysis. **d.** A Tet-On–inducible system was used to upregulate NeuroD2 in PNCs. Neurons were infected at DIV3 with a doxycycline-responsive NeuroD2 construct, allowing controlled induction before KCl stimulation. **e.** Immunostaining showing a strong increase in NeuroD2 protein following doxycycline treatment. Scale bar: 10 μm. **f.** Activity-repressed neuronal identity genes (*Hpca*, *Foxo6*, *Rtn4r*) remain downregulated following KCl stimulation despite NeuroD2 overexpression (n = 4; mean ± SEM; one-way ANOVA). **g-i.** Fos loss-of-function analysis. **g.** Lentiviral construct containing the dCas9-KRAB system and guide RNAs targeting the Fos promoter was designed to reduce its transcriptional induction following neuronal stimulation. **h.** Experimental workflow: PNCs were infected at DIV3, allowed to mature, and then exposed to KCl to induce activity-dependent Fos expression under conditions of targeted repression. **i.** Activity-repressed neuronal identity genes remain downregulated following KCl stimulation despite Fos knockdown, indicating that reducing activity-induced FOS alone is insufficient to prevent their repression (n = 4; mean ± SEM). **j-l.** Fos gain-of-function analysis. **j.** Tet-On lentiviral construct expressing human FOS was introduced into PNCs at DIV3; Fos was induced with doxycycline in the absence of KCl stimulation. **k.** Immunostaining confirms strong Fos overexpression (scale bar: 10 μm). **l.** FOS overexpression is sufficient to downregulate neuronal identity genes in the absence of neuronal stimulation (n = 4; mean ± SEM).

Conversely, CRISPR–KRAB-mediated silencing of *Fos* (**Fig. 7g-h** and **Supp. Fig. S5e**) did not interfere with IEG induction (**Supp. Fig. S5f**) or prevent the downregulation of neuronal identity genes (**Fig. 7i**), presumably due to redundancy among AP-1 subunits. However, strikingly, FOS overexpression in the absence of depolarization (**Fig. 7j-k** and **Supp. Fig. S5g**) was sufficient to repress all three neuronal identity genes tested (**Fig. 7l**), without broadly inducing the canonical activity-dependent transcriptional program (**Supp. Fig. S5h**). These results indicate that although FOS loss does not prevent repression—likely because of redundancy among AP-1 subunits—increased FOS expression is sufficient to reproduce the repression of neuronal identity genes observed during neuronal activation.

Overall, these results indicate that KAT3 shuttling accompanies both pathological and physiological neuronal activation and can be reproduced in cultured neurons. Manipulating NeuroD2 levels was insufficient to alter the activity-dependent repression of neuronal identity genes, consistent with redundancy within the identity-associated TF network. By contrast, although partial FOS depletion did not block this repression, FOS overexpression was sufficient to reproduce it in the absence of stimulation. Together, these findings identify increased FOS availability as a sufficient trigger for the identity-gene repression associated with neuronal activation and are consistent with a model in which AP-1 induction shifts the balance between competing KAT3-dependent transcriptional programs.

## Discussion

### Role of competitive KAT3 shuttling in plasticity

Neuronal activation induces profound changes in gene expression, most notably the rapid transcriptional burst that follows strong synaptic input and contributes critically to multiple forms of plasticity, including memory formation. However, much less is known about how activity-dependent transcription interacts with other transcriptional programs, particularly those that maintain neuronal identity. Our multi-omic analyses reveal that neuronal activation engages a competitive mechanism for the recruitment of the transcriptional coactivators CBP and p300. Specifically, activity-induced AP-1 factors and proneural terminal selectors such as NeuroD2 appear to compete for access to a limited pool of KAT3 proteins. Promoters and enhancers that gained KAT3 binding also showed increased chromatin accessibility, *de novo* AP-1 binding, and strengthened enhancer-promoter interactions. By contrast, enhancers— and, to a lesser extent, promoters—that lost KAT3 binding and H3K27ac displayed only modest reductions in accessibility and remained occupied by neuronal terminal selectors. Moreover, our functional assays in PNCs indicate that FOS expression is sufficient to reproduce the repression of neuronal identity genes observed during activation, even in the absence of stimulation. Together, these findings support a model in which AP-1, and possibly other activity-induced TFs, transiently redirect CBP/p300 away from neuronal identity programs toward plasticity-related regulatory elements.

Functionally, this competition is likely to have important implications for plasticity. Why might neurons employ a competitive mechanism that transiently challenges the maintenance of neuronal identity?

A likely explanation is that limiting CBP/p300 availability imposes temporal and cellular precision on activity-dependent transcription. Precise temporal control of gene expression is essential for memory formation and specificity. The pioneer-like activity of AP-1 (Su et al., 2017) together with the efficient recruitment of KAT3 proteins to AP-1 bound sites may ensure a rapid and robust transcriptional burst, thereby facilitating the molecular cascades that underlie long-term potentiation and memory consolidation. Competition for a limiting coactivator pool could therefore help prioritize strongly induced regulatory programs without requiring global increases in CBP/p300 abundance. By forcing competition between identity- and plasticity-associated transcription factors, neurons can generate a rapid but selective transcriptional burst in response to stimulation while restricting indiscriminate coactivator redistribution. In this view, transient redistribution of CBP/p300 is not simply a consequence of activation, but may contribute to sharpening plasticity responses without fully destabilizing neuronal identity. Consistent with this idea, studies of CBP-deficient mice showed that restricting CBP levels reduced the expression of specific pre- and postsynaptic proteins and decreased neuronal excitability, without appreciably compromising neuronal identity (Chen et al., 2010; Barrett et al., 2011; Valor et al., 2011; Lipinski et al., 2022). Conversely, excessive CBP/p300 availability might also perturb the balance between competing transcriptional programs. Consistent with the importance of dosage, duplication of the *CREBBP* gene in chromosome 16p13.3 duplication syndrome is also associated with intellectual disability (OMIM #613458), albeit generally less severe than that caused by haploinsufficiency of the same gene (RSTS1, OMIM #180849).

An additional, non-mutually exclusive possibility is that this redistribution limits transcriptional activity at neuronal identity loci during periods of intense activation. The transcriptional burst accompanying neuronal activation has been linked to DNA double-strand breaks that must be precisely repaired (Madabhushi et al., 2015). In this context, the transient reduction of CBP/p300 occupancy at large identity-associated regulatory domains could reduce transcriptional engagement at these loci during periods of exceptionally high activity. Whether this provides protection from activity-associated transcriptional stress remains to be tested

### Competitive KAT3 shuttling during development and aging

Activity-dependent redistribution of CBP/p300 may reflect the reuse of a regulatory logic also deployed during development. During neuronal differentiation, bHLH transcription factors help establish cell type-specific regulatory regions and long-range chromatin interactions (Bonev et al., 2017), whereas other developmental transitions are accompanied by increased expression of AP-1 components and repression of identity-associated factors (Stroud et al., 2020). Although these studies do not directly address KAT3 redistribution, they suggest that shifts in the balance between identity-maintaining and inducible transcription factor programs may be a recurring feature of neuronal state transitions.

Aging has also been linked to chromatin remodeling at cis-REs that balance identity and plasticity programs. A recent large-scale study analyzing age-related chromatin accessibility and matched transcriptional changes across 22 primary mouse cell types identified a consistent pattern: cis-REs that lost accessibility with age were enriched for cell-identity TF motifs, whereas cis-REs that gained accessibility were enriched for AP-1 motifs (Patrick et al., 2024). Based on these observations, the authors proposed the “Stimulus-Induced Programming Hijacks Ontogeny” (SIPHON) model, in which AP-1-driven chromatin opening, beneficial during maturation, becomes maladaptive when chronically re-engaged later in life by stress, inflammation, or other systemic signals, disrupting identity maintenance and accelerating aging. Supporting this framework, AP-1 factors have been implicated in maturation processes including neuronal differentiation (Herring et al., 2022), are broadly upregulated with age (Patrick et al., 2024), and include well-established “global aging genes” such as *Fos* and *Junb* (Almanzar et al., 2020; Zhang et al., 2021b).

Interestingly, neurons emerged as a notable exception to this general pattern. Whereas most cell types showed a roughly balanced number of cis-REs opening and closing with age, neurons displayed a strong predominance of closing regions and very little age-associated cis-RE opening. In addition, the coupling between differentially accessible regions and age-regulated gene expression was minimal in neurons compared with other cell types.

Our results suggest that neurons may experience a variant of the SIPHON mechanism. Rather than undergoing widespread AP-1-dependent opening of late-life cis-REs, neurons may respond to chronic AP-1 engagement by redistributing limiting CBP/p300 from identity-associated enhancers to stimulus-responsive loci. This would shift the balance between identity and plasticity programs without requiring extensive de novo chromatin opening and provides a potential framework for understanding the characteristic pattern of neuronal aging, in which identity erosion is more prominent than large-scale cis-RE activation.

At the same time, our previous work indicates that neuronal identity is remarkably robust: complete silencing of neuronal genes in mature neurons required simultaneous depletion of both CBP and p300, and even a single active allele out of the four combined *Crebbp*/*Ep300* alleles was sufficient to maintain identity programs (Lipinski et al., 2020). Large regulatory domains such as super-enhancers may contribute to this robustness by concentrating TFs and coactivators, thereby stabilizing transcriptional output (Whyte et al., 2013a; Sabari et al., 2018). Such robustness may explain how neurons tolerate repeated transient KAT3 redistribution during physiological activity without permanently compromising cell identity.

### Role of competitive KAT3 shuttling in disease

Although competitive KAT3 shuttling may contribute to normal plasticity, our analyses raise the possibility that persistent or excessive neuronal activation could chronically divert CBP/p300 away from identity-maintenance programs, thereby contributing to the neuronal identity erosion in pathological conditions characterized by sustained neuronal activity. Epilepsy represents an obvious candidate: experimental and clinical transcriptomic studies frequently report downregulation of neuronal and synaptic gene programs, a pattern consistent with competition between activity-induced and identity-maintenance transcriptional networks. However, many early studies relied on bulk RNA-seq, in which the apparent downregulation of neuronal identity genes could also reflect neuronal loss or gliosis.

A similar phenomenon may also occur in AD, where neuronal hyperactivity has been reported early in disease progression and is accompanied by broad disruption of neuronal transcriptional programs. Several transcriptomic studies show that neurons in AD partially downregulate neuronal-specific gene programs while upregulating stress, cell-cycle, or glial-associated programs. Single-nucleus analyses of human AD brain have revealed broad reductions in synaptic and neuronal gene expression across excitatory and inhibitory neuronal populations (Mathys et al., 2024). Grubman et al. reported altered neuronal regulatory programs in AD (Grubman et al., 2019), while Leng et al. identified transcriptional trajectories associated with progressive loss of neuronal subtype-specific features (Leng et al., 2021). Moreover, chromatin accessibility and multi-omic studies have identified losses of accessibility at neuronal regulatory elements linked to synaptic and identity-associated genes (Leng et al., 2021; Xiong et al., 2023). Together, these observations identify neuronal identity erosion as a central mechanism linking persistent neuronal activation to cognitive decline and neurological disease.

Notably, competition between viral and endogenous proteins for the CBP/p300 complex has been linked to disease in other biological contexts (Polansky and Schwab, 2020). In the nervous system, reduced CBP availability has been implicated in multiple disorders, from intellectual disability (Chen et al., 2010; Barrett et al., 2011; Valor et al., 2011; Lipinski et al., 2022) to neurodegeneration and neuronal identity erosion in Huntington’s disease (Giralt et al., 2012), among other examples. Our findings extend this concept by suggesting that CBP/p300 availability can also be dynamically redistributed between endogenous transcriptional programs in response to neuronal activity. Competition for CBP/p300 may be relevant in neuropathological states associated with persistent or chronic neuronal activation, such as epilepsy or Alzheimer’s disease, in which sustained redistribution of CBP/p300 away from identity-maintenance programs could favor maladaptive transcriptional states and progressively compromise neuronal function.

## Acknowledgments

We thank Ana Pombo, Christoph Thieme and Alexander Kukalev for their advice in GAM analysis, and Marc Marti-Renom and Leo Zuber for their support in METALoci analyses. We thank the personnel of the Mouse facility, and the Omics and microscopy core services at the Instituto de Neurociencias for their assistance. We also thank R. Olivares and C. Racovac for technical assistance. S.N. was recipient of a fellowship from the Spanish Ministry of Science and Innovation (MICINN). J.P-L. was the recipient of a CIAPOS contract given by Generalitat Valenciana and European Social Found. P.T-R. and M.C. are recipients of fellowships from Generalitat Valenciana. J.M.S-P. is funded by “Emergia” Grant EMC21_00188 by Junta de Andalucía; grant PID2023-147172NA-I00 by MICIU/AEI/ 10.13039/501100011033 and by the “European Union”; and grant CNS2023-143945 by MICIU/AEI/10.13039/501100011033 and by “European Union NextGenerationEU/PRTR.” A.B. research is supported by grants PID2023-148442NB-I00 by MICIU/AEI/10.13039/501100011033 co-financed by ERDF, FCAIXA HR22-00394 from Fundación LaCaixa, and CIPROM/2023/15 from the Generalitat Valenciana. The Instituto de Neurociencias is a “Centre of Excellence Severo Ochoa” (CEX2021-001165-S).

## Author contributions

Conceptualization, S.N. and A.B.; Methodology, M.C., B.dB., J.P-L., P.T-R., M.L. and F.M.; Software, S.N., and J-M.S-P.; Investigation, S.N., J.P-L., M.C., B.dB., P.T-R., and F.M.; Data Curation and Visualization, S.N.; Writing - Original Draft, S.N. and A.B.; Supervision, A.B.; Funding Acquisition, A.B.

## Competing interests

The authors declare no competing interests.

## Declaration of generative AI and AI-assisted technologies in the writing process

During the preparation of this work the authors used ChatGTP to revise English grammar and usage. After using this tool/service, the authors reviewed and edited the content as needed and take full responsibility for the content of the publication.

## Methods

### Mouse strains and treatments

The experiments were carried out using wild-type C57BL/6 J male and female mice between 2 and 5 months of age. For KA-induced neuronal activation, KA was administered intraperitoneally at 25 mg/kg (Milestone PharmTech USA Inc.), either as a single injection or divided into two equal doses administered 30 min apart (ChIP-seq experiments for NeuroD2 and H3K27ac). Animals were monitored following KA administration to ensure the development of stage 4 seizures before sample collection at 1 h. All mice were bred under specific pathogen-free (SPF) conditions at the Animal House of the Instituto de Neurociencias (CSIC-UMH), under a 12 h light/12 h dark cycle (7:00 a.m. to 7:00 p.m.) at 20–24 °C and 40–60% relative humidity, with ad libitum access to food and water. All animal protocols were performed in accordance with Spanish and European regulations and were approved by the Animal Welfare Committee of the Instituto de Neurociencias, the CSIC Ethical Committee, and the Dirección General de Agricultura, Ganadería y Pesca de la Generalitat Valenciana.

### Novelty exposure

For the novel environment (NE) experiment, a total of four 3-month-old female wild-type mice were individually placed for 60 min in a novel arena. The arena consisted of a white acrylic box (48 × 48 × 30 cm) containing objects of different shapes and colors. The mice were then processed for perfusion.

### Western blot

Two-month-old wild-type female mice were euthanized by cervical dislocation, and the hippocampi were microdissected, immediately frozen on dry ice, and stored at −80 °C. Samples were lysed with 300 µl Co-IP lysis buffer (50 mM Tris-HCl, 150 mM NaCl, 1% NP-40, 1% SDS, and 1 mM EDTA) and supplemented with protease inhibitors. Equal amounts of protein (80 µg) were mixed with 4× Laemmli sample buffer, boiled at 95 °C for 5 min, and separated by SDS-PAGE on 6.5% polyacrylamide gels. Proteins were transferred onto PVDF membranes (Millipore) using a wet transfer system overnight at 4 °C. Membranes were blocked with 5% non-fat dry milk in Tris-buffered saline containing 0.1% Tween-20 (TBST) for 2 h at room temperature and then incubated overnight at 4 °C with primary antibodies. The following primary antibodies were used: anti-CBP (D6C5) [#cs-7389, Cell Signaling; 1:1000]; anti-p300 (C-20) [#sc-585, Santa Cruz Biotechnology; 1:1000]; anti-NeuroD2 (C-20) [ab109406, Abcam; 1:1000]; and anti-Vinculin (42H89L44) [#700062, Invitrogen; 1:5000].The next day, the membranes were incubated with goat anti-rabbit IgG (H+L) HRP-conjugated secondary antibody [#31460, Invitrogen; 1:5000]. Protein bands were visualized using enhanced chemiluminescence (ECL) and imaged using a ChemiDoc system. Band intensities were quantified with ImageLab software and normalized to Vinculin.

### Primary hippocampal cultures and lentiviral infection

Primary hippocampal neuronal cultures were prepared from ICR embryos as previously described (Scandaglia et al., 2015). Briefly, hippocampi were dissected, pooled, and mechanically dissociated to obtain a neuronal cell suspension. Cells were plated in 24-well plates at a density of 0.13 × 10^6 neurons per well. At DIV3, cultures were infected with the indicated viral constructs. For functional assays, neurons were co-infected at DIV3 and maintained for 14 days before experimental processing, whereas cultures used for in vitro validation experiments were maintained for 19 days. To suppress spontaneous neuronal activity prior to stimulation, cultures were treated for 24 h with tetrodotoxin citrate (TTX; 1 µM) and D(-)-2-amino-5-phosphonopentanoic acid (D-AP5; 100 µM). Neuronal activity was subsequently induced by KCl-mediated depolarization. Briefly, pre-warmed KCl depolarization buffer containing 170 mM KCl, 2 mM CaCl₂, 1 mM MgCl₂, and 10 mM HEPES was added directly to the culture medium to achieve a final concentration of 31% (v/v) of depolarization buffer in each well. For immunohistochemistry experiments, neurons were cultured on poly-L-lysine-coated glass coverslips.

### Immunohistochemistry and microscopy

For immunocytochemistry, cells were fixed in 4% paraformaldehyde for 12 min at room temperature, washed with PBS, and permeabilized with PBS containing 0.2% Triton X-100 for 30 min at room temperature. Cells were incubated with primary antibodies overnight at 4 °C, and with secondary antibodies for 1–3 h at room temperature. Finally, nuclei were counterstained with DAPI (Invitrogen). Images were acquired using a Leica SPEII upright confocal microscope and processed using Fiji-ImageJ software with the *Cell Counter* plugin.

For immunohistochemistry, mice were anesthetized with a mixture of ketamine and xylazine in PBS, then perfused first with PBS to remove blood, followed by 4% paraformaldehyde (PFA) via injection into the left ventricle. Brains were carefully removed, incubated in 4% PFA overnight, and sectioned into 50 μm slices using a vibratome. Antigen retrieval was performed before immunostaining when necessary. For antigen retrieval, brain sections were incubated in 10 mM sodium citrate buffer with 0.05% Tween-20 (pH 6) at 80°C for 30 minutes, then allowed to cool to room temperature. After three 5-minute washes in PBS with 0.3% Triton X-100 (PBS-T), tissue slices were incubated in blocking solution (4% heat-inactivated newborn calf serum in PBS-T) for 2 hours at room temperature with gentle agitation. The primary antibodies were added at the appropriate concentrations and incubated overnight at 4°C with agitation. The primary antibodies used were: anti-CBP (C1) [sc-7300, Santa Cruz Biotechnology, 1:250]; anti-p300 (C-20) [sc-585 X, Santa Cruz Biotechnology, 1:100]; anti-FOS [#226 017 made in rat, Synaptic Systems, 1:1000]; anti-FOS [MA5-15055 made in mouse, Thermo scientific, 1:500]; α-NeuroD2 [ab109406, Abcam, 1:250]; and anti-Flag M2 [F1804, Sigma-Aldrich, 1:500]. Following primary antibody incubation, sections were rinsed three times for 5 minutes in PBS-T, then incubated with the secondary antibody for 1.5 hours at room temperature with agitation. Sections were again rinsed three times for 5 minutes in PBS-T. For DNA counterstaining, brain slices were incubated with 1 nM DAPI (4’,6-diamidino-2-phenylindole; Invitrogen) in PBS for 10 minutes at room temperature, followed by three 5-minute washes in PBS. Sections were mounted using Fluoromount aqueous mounting medium (Sigma) and sealed with nail polish. Confocal images were obtained using a vertical Confocal Microscope Leica SPEII with oil-immersion 40x objective lens (NA 1.4) and zoom 1.5, with a 1024×1024 collection box. Confocal pinhole was set to 1 AU for each channel and pixel/voxel size was 1µm. Images were processed as Maximum Intensity Projection with Fiji software (ImageJ 1.53f51) and the mean intensity was calculated to measure the immunoreactivity, relativized to DAPI. For super-resolution imaging analysis, tissue samples were imaged on a Zeiss LSM 880 confocal microscope with Airyscan and Elyra PS.1, using a 63x PlanApo oil-immersion objective (NA 1.4) and processed with ZEN 2.3 software (Zeiss, RRID: SCR_013672), maintaining identical laser power, photomultiplier gain, pinhole, and detection filter settings. Image size and voxel spacing were 1912 x 1912 pixels, with 16-bit depth and a voxel spacing of 0.18 µm. All super-resolution images were then deconvolved with Huygens Professional version 25.10 (Scientific Volume Imaging, The Netherlands, http://svi.nl). The colocalization analysis was made using automatic thresholding within the ‘Coloc’ tool in Imaris 10.2.0 software on the 3D image. Crosstalk between fluorophores was eliminated adjusting the detector spectral ranges and using sequential scanning for image acquisition.

### RNA isolation and RT-qPCR

Total RNA was isolated from hippocampal neuronal cultures using TRI Reagent (Sigma-Aldrich) according to the manufacturer’s instructions. Purified RNA was reverse transcribed into complementary DNA (cDNA) using the RevertAid First-Strand cDNA Synthesis Kit (Fermentas). Quantitative PCR was subsequently performed using EvaGreen-based qPCR reagents on either an Applied Biosystems 7300 Real-Time PCR System or a QuantStudio 3 Real-Time PCR System. Gene expression was analyzed using the primer pairs listed below.

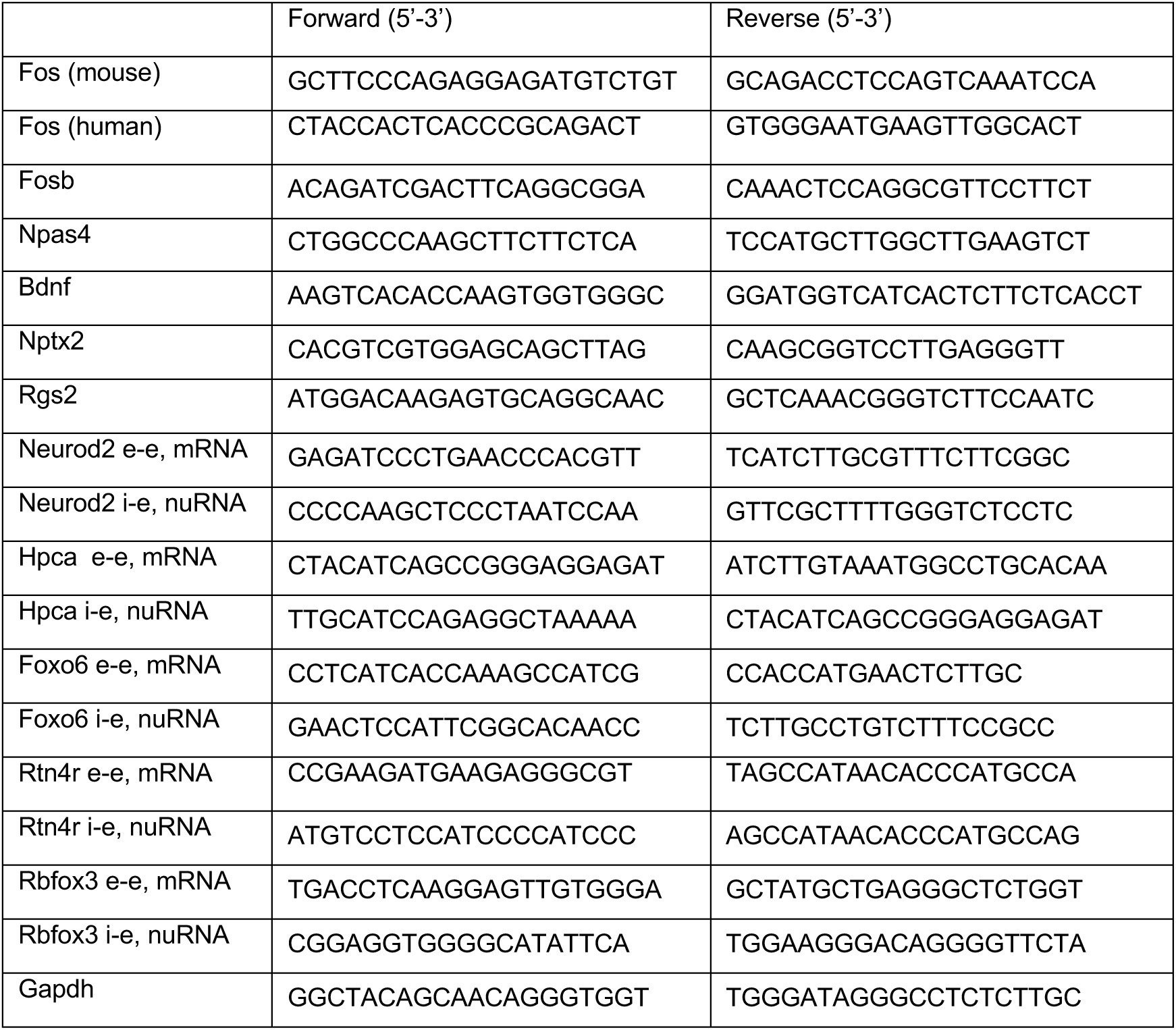

### RNA-seq experiments and analysis

For in vivo tissue samples, hippocampi were collected from 4-month-old male mice, including two biological replicates for condition. Mice were euthanized by cervical dislocation, the brain was immediately removed, and the hippocampi were dissected. For each replicate, hippocampi from three mice were pooled. Hippocampi were placed in SafeLock Eppendorf tubes together with TRI Reagent (Sigma-Aldrich) and zirconium oxide beads 0,5 mm of diameter (Next Advance, ZrOB05). Samples were homogenized in a Bullet blender Storm 24 for 3 minutes. For in vitro assays, primary hippocampal cultures were detached and directly homogenized using TRI Reagent (Sigma-Aldrich). For RNA isolation, Chloroform was added to each sample and vigorously mixed by vortex. After 3 min of incubation, samples were centrifuged for 15 min at 12,000 x g at 4 °C. The aqueous phase was transferred to a new Eppendorf tube and mixed with isopropanol. To facilitate nucleic acid precipitation, 10 μg of RNase-free glycogen was added to the samples. After 30 min at room temperature, samples were centrifuged for 1 h at 13,000 rpm at 4 °C. The supernatant was discarded, and the pellet was washed with 75% ethanol and was allowed to air-dry for 15 min. Finally, the pellet was resuspended in RNase-free water. To eliminate genomic DNA, samples were treated with RNase-free DNase I (Qiagen) for 30 min at 25 °C. RNA was precipitated with phenol-chloroform-isoamyl alcohol (Sigma-Aldrich). The supernatant was discarded, and the pellet was washed with 80% ethanol. The RNA pellet was air-dried for 15 min and dissolved in RNase-free water. RNA quality and concentration were assessed using a NanoDrop spectrophotometer (Thermo Fisher) and Bioanalyzer (Agilent). RNA was sequenced using polyA selection for in vivo tissue samples and the sequencing libraries for in vitro cultured neurons data were generated from ribosomal RNA-depleted RNA. For all RNA datasets, reads were aligned to GRCm38 using STAR (v2.6.1) (Dobin et al., 2013). SAMtools (v1.1) was used to retain reads with MAPQ > 30. For mRNA and riboRNA datasets, counts were quantified over exons, whereas nuRNA counts were quantified at the gene level using Rsubread (v2.20) and the Mus_musculus.GRCm38.99.gtf annotation from Ensembl. DESeq2 (v1.46) was used for differential expression analysis between saline (SAL) and KA 1 h conditions in the nuRNA, mRNA, and riboRNA datasets. ChromHMM (Ernst and Kellis, 2012) was used to define transcriptionally active regions for metagene analysis based on the presence of CBP, H3K27ac, and RNAP2 signals. These regions were separated into increased or decreased categories based on CBP differential binding and annotated to the nearest Ensembl gene ID. nuRNA bigWig tracks normalized by RPKM were generated using deepTools (Ramírez et al., 2014), and metagene profiles were generated using plotProfile. For primary hippocampal cultures, RNA-seq was performed on total RNA using paired-end sequencing, and the resulting data were processed using the same bioinformatic pipeline described above.

### ChIP-seq experiments and analysis

Chromatin immunoprecipitation (ChIP) experiments were performed using 2– 3-month-old wild-type (WT) male mice. CBP (Santa Cruz Biotechnology, sc-583) and p300 (Santa Cruz Biotechnology, sc-585) ChIP experiments were performed as previously described (Lipinski et al., 2020), whereas NEUROD2 (Abcam ab109406) and H3K27ac (Abcam ab4729) ChIP experiments were performed following the protocol described by (Alaiz-Noya et al., 2025). The FOS ChIP experiment (Thermo Fisher Scientific #MA5-15055, clone T.142.5) was also performed as described by (Alaiz-Noya et al., 2025). For CBP and p300 ChIP experiments, biological replicates consisted of n = 2 saline-treated and n = 3 KA-treated mice. For NEUROD2 and H3K27ac ChIP experiments, n = 3 biological replicates were analyzed per condition (saline and KA). ChIP-seq reads were trimmed using Trim Galore (v0.6.4) and aligned with Bowtie2 (v2.3.4.3) (Langmead and Salzberg, 2012). Blacklist regions were removed, and only reads mapping to nuclear chromosomes with MAPQ > 30 were retained for further analysis using SAMtools (v1.1) (Li et al., 2009). Peak calling was performed with MACS2 (v2.1.1.20160306) (Zhang et al., 2008), and read counts were extracted and differential binding regions were identified using DiffBind (v3.16) (Ross-Innes et al., 2012). ChIPpeakAnno (v3.20.1) (Zhu et al., 2010) was used to annotate regions to genes and genomic positions. Promoters were defined as regions annotated as promoters; intragenic regions included exons, introns, and 5’ and 3’ UTRs; and intergenic regions comprised all remaining annotations. Nearest-neighbor CBP category analysis was performed using the *closest* command in bedtools (v2.30.0) (Quinlan and Hall, 2010). Signal quantification was performed using either HOMER tags or RPGC-normalized bigWig files (rtracklayer v1.66.0; GenomicRanges v1.58.0) (Lawrence et al., 2009; Lawrence et al., 2013). Read counts were also obtained directly from scaled ChIP-seq BAM files for calculation of log2 fold changes. Overlap between TF peaks for the UpSet plot (Conway et al., 2017) was calculated with ChIPpeakAnno using findOverlapsOfPeaks. For the classification shown in **Figures 1g and 3d**, CBP-bound regions were grouped into ten categories based on 0.5-log2-fold-change intervals, spanning values from < −1.5 to > 2.5, and signal from different marks was obtained using deepTools or annotated using ChIPpeakAnno. Motif analysis was performed with HOMER (v5.1) using findMotifsGenome.pl; for the enrichment analysis in **Supplementary Figure S3a**, createHomer2EnrichmentTable.pl was used.

### HiChIP H3K27ac

HiChIP for H3K27ac was performed as previously described (Di Giammartino et al., 2022), with modifications. Each biological replicate comprised six hippocampi from 3–5-month-old male mice. Hippocampi were cross-linked with 1% PFA, quenched with glycine, lysed, and nuclei were digested with MboI. Restriction fragment ends were filled in with biotinylated nucleotides and subjected to proximity ligation. Nuclei were then lysed and sonicated, and chromatin was immunoprecipitated overnight at 4 °C with an anti-H3K27ac antibody (Abcam ab4729, 2 µg per million nuclei). After extensive washing and elution, cross-links were reversed and DNA was purified. Biotinylated ligation products were captured with streptavidin beads, tagmented with TDE1 transposase, PCR-amplified, size-selected using AMPure XP beads, and sequenced. Library quality was assessed throughout the protocol by monitoring restriction digestion, ligation, sonication, H3K27ac enrichment, DNA concentration, and final library size distribution.

### Hi-C and HiChIP analysis

Hi-C and HiChIP datasets were analyzed as described by (Franke et al., 2021) with minimal modifications. Briefly, paired-end reads were aligned using BWA (v0.7.17) (Li and Durbin, 2009). Pairtools (v0.3.0) (Abdennur et al., 2024) was used to select valid pairs, retaining those where both ends mapped to a unique genomic position (UU) or one end was rescued from a chimeric read (UR or RU), ensuring confident positional assignment for both loci of each contact. Pairs were then sorted and deduplicated using pairtools sort and dedup. MboI restriction enzyme sites were computed using Cooler (v0.10.4) (Abdennur and Mirny, 2020), and dangling-end and self-circle pairs were subsequently filtered with pairtools. Valid pairs were converted to .hic format and binned into contact matrices using Juicer (v1.6) (Durand et al., 2016). Conversion to .cool format was performed with hic2cool, and contact pileups were generated with coolpuppy (v1.1.0) (Flyamer et al., 2020). For HiChIP, loop calling and differential loop analysis were performed with FitHiChIP (v11) (Bhattacharyya et al., 2019) using two biological replicates per condition for H3K27ac HiChIP in SAL and KA 1 h samples. Contact matrices were visualized using Juicebox, and genomic profiles and loops were visualized in IGV. To assess the association between TSSs and putative enhancers, the full list of loops was intersected with a BED file containing TSS genomic positions using bedtools pairtobed. The number of loops anchored at CBP-increased and CBP-reduced peaks was then quantified, and odds ratios were calculated to determine whether the observed associations deviated from chance and in which direction. For METALoci, the analysis was conducted as described by (Mota-Gómez et al., 2026). Genomic windows of 10 kb were used for chromatin contact data. For gene-transition analyses, vectors between SAL and KA conditions were calculated after obtaining local Moran’s I (LMI) classifications using the `lm -i` command.

### Genome architecture mapping (GAM) analysis

GAM data were analyzed and filtered as described by (Winick-Ng et al., 2021), with the exception that 20-kb genomic windows were used. NPMI matrices were computed, and the top 10% strongest contacts were selected. To assess co-occupancy of CBP and NeuroD2 at contact anchors, peaks from each factor were independently called and then intersected with contact anchors. Contacts were classified according to whether each factor was present at both anchors, at only the left or right anchor, or at neither anchor. For the permutation test, NeuroD2 peaks were shifted along each chromosome 10,000 times as a block, preserving the relative distances between peaks within the same chromosome. The observed/expected (Obs/Exp) enrichment ratio was calculated by dividing the observed co-occupancy count for each category by the mean count obtained across the 10,000 permutations.

### ATAC-seq analysis

ATAC-seq reads were trimmed using Trim Galore (v0.6.4) and aligned to the mouse reference genome mm10 using Bowtie2 (v2.3.5.1) in paired-end mode. Reads were filtered for MAPQ ≥ 30 and alignment to nuclear chromosomes using SAMtools (v1.1), blacklist regions were removed, and duplicate reads were removed using Picard (v2.27). Peak calling was performed with MACS2 (v2.1.1.20160306) using the --format BAMPE. Differential accessibility analysis was performed with DiffBind (v3.16). For footprint analysis, RGT-HINT ATAC (v0.12.3) (Li et al., 2019) was applied to CBP-defined regions. Coverage tracks were normalized by RPGC using deepTools (v3.5.0).

### Other bioinformatic analyses

Quality control of all ChIP-seq, RNA-seq and ATAC-seq data was performed with FastQC (v0.11.8), and adapter trimming was performed with Trim Galore (v0.6.4). Aligned files were processed with SAMtools, bedtools, and deepTools. Genomic data were visualized with IGV (v2.6.3 or v2.17.4) (Thorvaldsdóttir et al., 2013). Gene ontology enrichment analyses for CBP and H3K27ac datasets were performed with PANTHER (Thomas et al., 2022). Super-enhancers were identified using ROSE (Whyte et al., 2013b) from basal CBP and H3K27ac peaks. UCSC liftOver (Casper et al., 2026) was used to map the human H3K27ac-defined regions in BI_Brain_Hippocampus_Middle.bed to the mouse genome. Other analyses were performed using custom scripts in R v4.4.2, and plots were generated with ggplot2 (v4.0.1) (Wickham, 2016).

### Statistical analysis

Statistical analyses were performed using GraphPad Prism (v9.4.1; GraphPad Software, La Jolla, CA, USA) and RStudio (v4.5.3). For comparisons between two independent groups, data were first assessed for normality using the Shapiro–Wilk test. Normally distributed data were analyzed using a two-tailed unpaired Student’s t-test. For comparisons involving more than two groups, a one-way analysis of variance (ANOVA) was performed separately for each gene, followed by Bonferroni’s multiple-comparisons test for post hoc pairwise comparisons. For comparison of gene-expression distributions across conditions, data were standardized as row-wise Z-scores and analyzed using the Kruskal–Wallis test, followed by pairwise Wilcoxon rank-sum tests with Benjamini–Hochberg correction for multiple comparisons. Statistical significance was defined as p < 0.05.

### Data availability

The genomic datasets generated in this study will be deposited in the Gene Expression Omnibus (GEO) repository upon acceptance of the manuscript.

## Notes

### Competing Interest Statement

The authors have declared no competing interest.

## References

Abdennur N, Fudenberg G, Flyamer IM, Galitsyna AA, Goloborodko A, Imakaev M, Venev S V. (2024) Pairtools: From sequencing data to chromosome contacts. PLoS Comput Biol.

Abdennur N, Mirny LA (2020) Cooler: Scalable storage for Hi-C data and other genomically labeled arrays. Bioinformatics.

Achour M, Le Gras S, Keime C, Parmentier F, Lejeune FX, Boutillier AL, Neri C, Davidson I, Merienne K (2015) Neuronal identity genes regulated by super-enhancers are preferentially down-regulated in the striatum of Huntington’s disease mice. Hum Mol Genet 24:3481–3496 Available at: http://www.ncbi.nlm.nih.gov/pubmed/25784504.

Alaiz-Noya M, Miozzo F, Fuentes-Ramos M, Machnicka MA, Kurowska M, Herrera ML, del Blanco B, Ninerola S, Bustos-Martínez I, Wilczynski B, Barco A (2025) Neuronal type-specific modulation of cognition and AP-1 signaling by early-life rearing conditions. Nat Commun 16:9710 Available at: https://www.nature.com/articles/s41467-025-65343-5 [Accessed February 7, 2026].

Alarcon JM, Malleret G, Touzani K, Vronskaya S, Ishii S, Kandel ER, Barco A (2004) Chromatin acetylation, memory, and LTP are impaired in CBP+/- mice: a model for the cognitive deficit in Rubinstein-Taybi syndrome and its amelioration. Neuron 42:947–959 Available at: http://www.ncbi.nlm.nih.gov/entrez/query.fcgi?cmd=Retrieve&db=PubMed&dopt=Citation&list_uids=15207239.

Almanzar N et al. (2020) A single-cell transcriptomic atlas characterizes ageing tissues in the mouse. Nature 2020 583:7817 583:590–595.

Barrett RM, Malvaez M, Kramar E, Matheos DP, Arrizon A, Cabrera SM, Lynch G, Greene RW, Wood MA (2011) Hippocampal focal knockout of CBP affects specific histone modifications, long-term potentiation, and long-term memory. Neuropsychopharmacology 36:1545–1556.

Beagan JA, Pastuzyn ED, Fernandez LR, Guo MH, Feng K, Titus KR, Chandrashekar H, Shepherd JD, Phillips-Cremins JE (2020) Three-dimensional genome restructuring across timescales of activity-induced neuronal gene expression. Nat Neurosci 23:707– 717.

Beagrie RA, Scialdone A, Schueler M, Kraemer DCA, Chotalia M, Xie SQ, Barbieri M, De Santiago I, Lavitas LM, Branco MR, Fraser J, Dostie J, Game L, Dillon N, Edwards PAW, Nicodemi M, Pombo A (2017) Complex multi-enhancer contacts captured by genome architecture mapping. Nature 543:519–524.

Benito E, Barco A (2015) The Neuronal Activity-Driven Transcriptome. Mol Neurobiol 51:1071– 1088.

Bhattacharyya S, Chandra V, Vijayanand P, Ay F (2019) Identification of significant chromatin contacts from HiChIP data by FitHiChIP. Nat Commun.

Bonev B, Mendelson Cohen N, Szabo Q, Fritsch L, Papadopoulos GL, Lubling Y, Xu X, Lv X, Hugnot JP, Tanay A, Cavalli G (2017) Multiscale 3D Genome Rewiring during Mouse Neural Development. Cell 171:557–572 e24 Available at: https://www.ncbi.nlm.nih.gov/pubmed/29053968.

Calderon L, Weiss FD, Beagan JA, Oliveira MS, Georgieva R, Wang YF, Carroll T, Dharmalingam G, Gong W, Tossell K, de Paola V, Whilding C, Ungless MA, Fisher AG, Phillips-Cremins JE, Merkenschlager M (2022) Cohesin-dependence of neuronal gene expression relates to chromatin loop length. Elife 11 Available at: https://pubmed.ncbi.nlm.nih.gov/35471149/ [Accessed April 9, 2026].

Casper J et al. (2026) The UCSC Genome Browser database: 2026 update. Nucleic Acids Res.

Chen G, Zou X, Watanabe H, van Deursen JM, Shen J (2010) CREB binding protein is required for both short-term and long-term memory formation. J Neurosci 30:13066–13077.

Conway JR, Lex A, Gehlenborg N (2017) UpSetR: An R package for the visualization of intersecting sets and their properties. Bioinformatics.

Creyghton MP, Cheng AW, Welstead GG, Kooistra T, Carey BW, Steine EJ, Hanna J, Lodato MA, Frampton GM, Sharp PA, Boyer LA, Young RA, Jaenisch R (2010) Histone H3K27ac separates active from poised enhancers and predicts developmental state. Proc Natl Acad Sci U S A 107:21931–21936.

Cummings DM, Benway TA, Ho H, Tedoldi A, Fernandes Freitas MM, Shahab L, Murray CE, Richard-Loendt A, Brandner S, Lashley T, Salih DA, Edwards FA (2017) Neuronal and Peripheral Pentraxins Modify Glutamate Release and may Interact in Blood-Brain Barrier Failure. Cereb Cortex 27:3437–3448 Available at: https://pubmed.ncbi.nlm.nih.gov/28334103/ [Accessed April 5, 2026].

del Blanco B, Niñerola S, Martín-González AM, Paraíso-Luna J, Kim M, Muñoz-Viana R, Racovac C, Sanchez-Mut J V., Ruan Y, Barco Á, Ninerola S, M. M-GA, Paraíso-Luna J, Kim M, Muñoz-Viana R, Racovac C, Sánchez-Mut J V., Ruan Y, Barco Á (2024) KDM1A safeguards the topological boundaries of PRC2-repressed genes and prevents aging-related euchromatinization in neurons. Nat Commun 15:1781 Available at: https://www.nature.com/articles/s41467-024-45773-3.

Deneris ES, Hobert O (2014) Maintenance of postmitotic neuronal cell identity. Nat Neurosci 17:899–907.

Di Giammartino DC, Polyzos A, Apostolou E (2022) Assessing Specific Networks of Chromatin Interactions with HiChIP. In: Methods in Molecular Biology.

Durand NC, Shamim MS, Machol I, Rao SSP, Huntley MH, Lander ES, Aiden EL (2016) Juicer Provides a One-Click System for Analyzing Loop-Resolution Hi-C Experiments. Cell Syst.

Dyson HJ, Wright PE (2016) Role of Intrinsic Protein Disorder in the Function and Interactions of the Transcriptional Coactivators CREB-binding Protein (CBP) and p300. J Biol Chem 291:6714–6722 Available at: https://www.ncbi.nlm.nih.gov/pubmed/26851278.

Ernst J, Kellis M (2012) ChromHMM: Automating chromatin-state discovery and characterization. Nat Methods.

Fang F, Xu Y, Chew KK, Chen X, Ng HH, Matsudaira P (2014) Coactivators p300 and CBP maintain the identity of mouse embryonic stem cells by mediating long-range chromatin structure. Stem Cells 32:1805–1816.

Fernandez-Albert J, Lipinski M, Lopez-Cascales MT, Rowley MJ, Martin-Gonzalez AM, del Blanco B, Corces VG, Barco A (2019) Immediate and deferred epigenomic signatures of in vivo neuronal activation in mouse hippocampus. Nat Neurosci 22:1718–1730.

Flyamer IM, Illingworth RS, Bickmore WA (2020) Coolpup.py: Versatile pile-up analysis of Hi-C data. Bioinformatics.

Franke M, De la Calle-Mustienes E, Neto A, Almuedo-Castillo M, Irastorza-Azcarate I, Acemel RD, Tena JJ, Santos-Pereira JM, Gómez-Skarmeta JL (2021) CTCF knockout in zebrafish induces alterations in regulatory landscapes and developmental gene expression. Nat Commun.

Giralt A, Puigdellivol M, Carreton O, Paoletti P, Valero J, Parra-Damas A, Saura CA, Alberch J, Gines S (2012) Long-term memory deficits in Huntington’s disease are associated with reduced CBP histone acetylase activity. Hum Mol Genet 21:1203–1216 Available at: http://www.ncbi.nlm.nih.gov/pubmed/22116937.

Goodman RH, Smolik S (2000) CBP/p300 in cell growth, transformation, and development. Genes Dev 14:1553–1577 Available at: http://www.ncbi.nlm.nih.gov/pubmed/10887150.

Gräff J, Tsai LH (2013) Histone acetylation: molecular mnemonics on the chromatin. Nat Rev Neurosci 14:97–111 Available at: http://www.ncbi.nlm.nih.gov/pubmed/23324667.

Griffith EC, West AE, Greenberg ME (2024) Neuronal enhancers fine-tune adaptive circuit plasticity. Neuron 112:3043–3057.

Grubman A, Chew G, Ouyang JF, Sun G, Choo XY, McLean C, Simmons RK, Buckberry S, Vargas-Landin DB, Poppe D, Pflueger J, Lister R, Rackham OJL, Petretto E, Polo JM (2019) A single-cell atlas of entorhinal cortex from individuals with Alzheimer’s disease reveals cell-type-specific gene expression regulation. Nat Neurosci 22:2087–2097 Available at: https://pubmed.ncbi.nlm.nih.gov/31768052/ [Accessed April 9, 2026].

Heinz S, Benner C, Spann N, Bertolino E, Lin YC, Laslo P, Cheng JX, Murre C, Singh H, Glass CK (2010) Simple combinations of lineage-determining transcription factors prime cis-regulatory elements required for macrophage and B cell identities. Mol Cell 38:576–589.

Hennig AK, Peng GH, Chen S (2013) Transcription coactivators p300 and CBP are necessary for photoreceptor-specific chromatin organization and gene expression. PLoS One 8:e69721 Available at: http://www.ncbi.nlm.nih.gov/pubmed/23922782.

Herring CA et al. (2022) Human prefrontal cortex gene regulatory dynamics from gestation to adulthood at single-cell resolution. Cell 185:4428–4447.e28.

Hnisz D, Abraham BJ, Lee TI, Lau A, Saint-André V, Sigova AA, Hoke HA, Young RA (2013) Super-enhancers in the control of cell identity and disease. Cell 155:934 Available at: https://pubmed.ncbi.nlm.nih.gov/24119843/ [Accessed April 9, 2026].

Jin T, Yang Y, Guo Y, Zhang Y, Le Q, Huang N, Liu X, Yu J, Ma L, Wang F (2025) Disturbed engram network caused by NPTX downregulation underlies aging-related contextual fear memory deficits. Cell Research 2025 35:9 35:656–674 Available at: https://www.nature.com/articles/s41422-025-01157-w [Accessed April 5, 2026].

Korzus E, Rosenfeld MG, Mayford M (2004) CBP histone acetyltransferase activity is a critical component of memory consolidation. Neuron 42:961–972 Available at: http://www.ncbi.nlm.nih.gov/entrez/query.fcgi?cmd=Retrieve&db=PubMed&dopt=Citation&list_uids=15207240.

Kwok RP, Lundblad JR, Chrivia JC, Richards JP, Bachinger HP, Brennan RG, Roberts SG, Green MR, Goodman RH (1994) Nuclear protein CBP is a coactivator for the transcription factor CREB. Nature 370:223–6.

Langmead B, Salzberg SL (2012) Fast gapped-read alignment with Bowtie 2. Nat Methods 9:357–359.

Lawrence M, Gentleman R, Carey V (2009) rtracklayer: An R package for interfacing with genome browsers. Bioinformatics.

Lawrence M, Huber W, Pagès H, Aboyoun P, Carlson M, Gentleman R, Morgan MT, Carey VJ (2013) Software for Computing and Annotating Genomic Ranges. PLoS Comput Biol.

Lee PR, Fields RD (2021) Activity-Dependent Gene Expression in Neurons. Neuroscientist 27:355–366 Available at: https://pubmed.ncbi.nlm.nih.gov/32727285/ [Accessed August 26, 2026].

Leng K, Li E, Eser R, Piergies A, Sit R, Tan M, Neff N, Li SH, Rodriguez RD, Suemoto CK, Leite REP, Ehrenberg AJ, Pasqualucci CA, Seeley WW, Spina S, Heinsen H, Grinberg LT, Kampmann M (2021) Molecular characterization of selectively vulnerable neurons in Alzheimer’s disease. Nat Neurosci 24:276–287 Available at: https://pubmed.ncbi.nlm.nih.gov/33432193/ [Accessed April 9, 2026].

Li H, Durbin R (2009) Fast and accurate short read alignment with Burrows-Wheeler transform. Bioinformatics.

Li H, Handsaker B, Wysoker A, Fennell T, Ruan J, Homer N, Marth G, Abecasis G, Durbin R (2009) The Sequence Alignment/Map format and SAMtools. Bioinformatics 25:2078– 2079.

Li Z, Schulz MH, Look T, Begemann M, Zenke M, Costa IG (2019) Identification of transcription factor binding sites using ATAC-seq. Genome Biol.

Lipinski M, del Blanco B, Barco A (2019) CBP/p300 in brain development and plasticity: disentangling the KAT’s cradle. Curr Opin Neurobiol 59:1–8 Available at: https://www.ncbi.nlm.nih.gov/pubmed/30856481.

Lipinski M, Muñoz-Viana R, del Blanco B, Marquez-Galera A, Medrano-Relinque J, Caramés JM, Szczepankiewicz AA, Fernandez-Albert J, Navarrón CM, Olivares R, Wilczyński GM, Canals S, Lopez-Atalaya JP, Barco A (2020) KAT3-dependent acetylation of cell type-specific genes maintains neuronal identity in the adult mouse brain. Nat Commun 11:2588 Available at: http://biorxiv.org/content/early/2019/11/12/836981.abstract.

Lipinski M, Niñerola S, Fuentes-Ramos M, Valor LM, del Blanco B, López-Atalaya JP, Barco A (2022) CBP Is Required for Establishing Adaptive Gene Programs in the Adult Mouse Brain. The Journal of Neuroscience 42:7984–8001.

Lopez-Atalaya JP, Valor LM, Barco A (2014) Epigenetic Factors in Intellectual Disability. In: Progress in Molecular Biology and Translational Science, pp 139–176 Available at: https://linkinghub.elsevier.com/retrieve/pii/B9780128009772000061.

Loven J, Hoke HA, Lin CY, Lau A, Orlando DA, Vakoc CR, Bradner JE, Lee TI, Young RA (2013) Selective inhibition of tumor oncogenes by disruption of super-enhancers. Cell 153:320–334 Available at: http://www.ncbi.nlm.nih.gov/pubmed/23582323.

Lundblad JR, Kwok RPS, Laurance ME, Harter ML, Goodman RH (1995) Adenoviral ElA-associated protein p300 as a functional homologue of the transcriptional co-activator CBP. Nature 374:85–88.

Madabhushi R, Gao F, Pfenning AR, Pan L, Yamakawa S, Seo J, Rueda R, Phan TX, Yamakawa H, Pao P-C, Stott RT, Gjoneska E, Nott A, Cho S, Kellis M, Tsai L-H (2015) Activity-Induced DNA Breaks Govern the Expression of Neuronal Early-Response Genes. Cell 161:1592–1605 Available at: http://www.cell.com/article/S0092867415006224/fulltext.

Malik AN, Vierbuchen T, Hemberg M, Rubin AA, Ling E, Couch CH, Stroud H, Spiegel I, Farh KK-H, Harmin DA, Greenberg ME (2014) Genome-wide identification and characterization of functional neuronal activity-dependent enhancers. Nat Neurosci 17:1330–1339 Available at: 10.1038/nn.3808.

Mathys H et al. (2024) Single-cell multiregion dissection of Alzheimer’s disease. Nature 632:858–868 Available at: https://pubmed.ncbi.nlm.nih.gov/39048816/ [Accessed April 9, 2026].

Mota-Gómez I, Rodríguez JA, Dupont S, Hurtado A, Cadenas V, Zuber L, Maceda I, Lao O, Jedamzick J, Kühn R, Lacadie S, García-Moreno SA, Torres M, Real FM, Acemel RD, Capel B, Marti-Renom MA, Lupiáñez DG (2026) Chromatin spatial analysis by METALoci unveils sex-determining 3D regulatory hubs. Nature Structural & Molecular Biology 2026 33:4 33:577–589 Available at: https://www.nature.com/articles/s41594-026-01749-z [Accessed August 25, 2026].

Nagy Z, Tora L (2007) Distinct GCN5/PCAF-containing complexes function as co-activators and are involved in transcription factor and global histone acetylation. Oncogene 26:5341–5357 Available at: https://pubmed.ncbi.nlm.nih.gov/17694077/ [Accessed August 13, 2026].

Nativio R, Lan Y, Donahue G, Sidoli S, Berson A, Srinivasan AR, Shcherbakova O, Amlie-Wolf A, Nie J, Cui X, He C, Wang LS, Garcia BA, Trojanowski JQ, Bonini NM, Berger SL (2020) An integrated multi-omics approach identifies epigenetic alterations associated with Alzheimer’s disease. Nature Genetics 2020 52:10 52:1024–1035 Available at: https://www.nature.com/articles/s41588-020-0696-0 [Accessed August 26, 2026].

Patrick R et al. (2024) The activity of early-life gene regulatory elements is hijacked in aging through pervasive AP-1-linked chromatin opening. Cell Metab 36:1858–1881.e23.

Patterson SL, Abel T, Deuel TA, Martin KC, Rose JC, Kandel ER (1996) Recombinant BDNF rescues deficits in basal synaptic transmission and hippocampal LTP in BDNF knockout mice. Neuron 16:1137–1145.

Pentz ES, Cordaillat M, Carretero OA, Tucker AE, Sequeira Lopez MLS, Ariel Gomez R (2012) Histone acetyl transferases CBP and p300 are necessary for maintenance of renin cell identity and transformation of smooth muscle cells to the renin phenotype. Am J Physiol Heart Circ Physiol 302:2545–2552.

Polansky H, Schwab H (2020) Dynamics of sequestering the limiting p300/CBP, viral cis-regulatory elements, and disease. J Biosci:79 Available at: http://www.ias.ac.in/jbiosci [Accessed April 7, 2026].

Quinlan AR, Hall IM (2010) BEDTools: A flexible suite of utilities for comparing genomic features. Bioinformatics.

Raisner R, Kharbanda S, Jin L, Jeng E, Chan E, Merchant M, Haverty PM, Bainer R, Cheung T, Arnott D, Flynn EM, Romero FA, Magnuson S, Gascoigne KE (2018) Enhancer Activity Requires CBP/P300 Bromodomain-Dependent Histone H3K27 Acetylation. Cell Rep 24:1722–1729 Available at: https://www.ncbi.nlm.nih.gov/pubmed/30110629.

Ramírez F, Dündar F, Diehl S, Grüning BA, Manke T (2014) DeepTools: A flexible platform for exploring deep-sequencing data. Nucleic Acids Res.

Ross-Innes CS, Stark R, Teschendorff AE, Holmes KA, Ali HR, Dunning MJ, Brown GD, Gojis O, Ellis IO, Green AR, Ali S, Chin SF, Palmieri C, Caldas C, Carroll JS (2012) Differential oestrogen receptor binding is associated with clinical outcome in breast cancer. Nature 481:389–393.

Sabari BR et al. (2018) Coactivator condensation at super-enhancers links phase separation and gene control. Science (1979) 361 Available at: https://www.ncbi.nlm.nih.gov/pubmed/29930091.

Saha RN, Wissink EM, Bailey ER, Zhao M, Fargo DC, Hwang JY, Daigle KR, Fenn JD, Adelman K, Dudek SM (2011) Rapid activity-induced transcription of Arc and other IEGs relies on poised RNA polymerase II. Nat Neurosci 14:848–856 Available at: https://www.ncbi.nlm.nih.gov/pubmed/21623364.

Shen Y et al. (2012) A map of the cis-regulatory sequences in the mouse genome. Nature 488:116–120 Available at: http://www.ncbi.nlm.nih.gov/pubmed/22763441.

Signal B, Phipps AJ, Giles KA, Huskins SN, Mercer TR, Robinson MD, Woodhouse A, Taberlay PC (2024) Ageing-Related Changes to H3K4me3, H3K27ac, and H3K27me3 in Purified Mouse Neurons. Available at: 10.3390/cells13161393.

Stroud H, Yang MG, Tsitohay YN, Davis CP, Sherman MA, Hrvatin S, Ling E, Greenberg ME (2020) An Activity-Mediated Transition in Transcription in Early Postnatal Neurons. Neuron 107:874–890.e8 Available at: https://linkinghub.elsevier.com/retrieve/pii/S0896627320304372.

Su Y, Shin J, Zhong C, Wang S, Roychowdhury P, Lim J, Kim D, Ming GL, Song H (2017) Neuronal activity modifies the chromatin accessibility landscape in the adult brain. Nat Neurosci 20:476–483 Available at: https://www.ncbi.nlm.nih.gov/pubmed/28166220.

Sugo N, Atsumi Y, Yamamoto N (2025) Transcription and epigenetic factor dynamics in neuronal activity-dependent gene regulation. Trends in Genetics 41:425–436.

Svensson K, LaBarge SA, Sathe A, Martins VF, Tahvilian S, Cunliffe JM, Sasik R, Mahata SK, Meyer GA, Philp A, David LL, Ward SR, McCurdy CE, Aslan JE, Schenk S (2020) p300 and cAMP response element-binding protein-binding protein in skeletal muscle homeostasis, contractile function, and survival. J Cachexia Sarcopenia Muscle 11:464– 477.

Telese F, Ma Q, Perez PM, Notani D, Oh S, Li W, Comoletti D, Ohgi KA, Taylor H, Rosenfeld MG (2015) LRP8-Reelin-Regulated Neuronal Enhancer Signature Underlying Learning and Memory Formation. Neuron 86:696–710 Available at: http://www.cell.com/article/S0896627315002573/fulltext.

Thomas PD, Ebert D, Muruganujan A, Mushayahama T, Albou LP, Mi H (2022) PANTHER: Making genome-scale phylogenetics accessible to all. Protein Sci 31:8–22 Available at: https://pubmed.ncbi.nlm.nih.gov/34717010/ [Accessed August 27, 2026].

Thorvaldsdóttir H, Robinson JT, Mesirov JP (2013) Integrative Genomics Viewer (IGV): High-performance genomics data visualization and exploration. Brief Bioinform 14:178–192.

Tutukova S, Tarabykin V, Hernandez-Miranda LR (2021) The Role of Neurod Genes in Brain Development, Function, and Disease. Front Mol Neurosci.

Valor LM, Pulopulos MM, Jimenez-Minchan M, Olivares R, Lutz B, Barco A (2011) Ablation of CBP in Forebrain Principal Neurons Causes Modest Memory and Transcriptional Defects and a Dramatic Reduction of Histone Acetylation But Does Not Affect Cell Viability. Journal of Neuroscience 31:1652–1663.

Wang S, Li T, Sheng C, Tan J, Yang Y, Ma X, Liu Y, Wei R, Zhou F, Zhou L, Wang X (2026) CBP/p300 is critical for the expansion and maintenance of functional pancreatic α cell mass. Nat Commun 17:5087 Available at: https://www.nature.com/articles/s41467-026-71499-5 [Accessed July 14, 2026].

Weinert BT, Narita T, Satpathy S, Srinivasan B, Hansen BK, Scholz C, Hamilton WB, Zucconi BE, Wang WW, Liu WR, Brickman JM, Kesicki EA, Lai A, Bromberg KD, Cole PA, Choudhary C (2018) Time-Resolved Analysis Reveals Rapid Dynamics and Broad Scope of the CBP/p300 Acetylome. Cell 174:231–244 e12.

Whyte WA, Orlando DA, Hnisz D, Abraham BJ, Lin CY, Kagey MH, Rahl PB, Lee TI, Young RA (2013a) Master transcription factors and mediator establish super-enhancers at key cell identity genes. Cell 153:307–319 Available at: https://www.ncbi.nlm.nih.gov/pubmed/23582322.

Whyte WA, Orlando DA, Hnisz D, Abraham BJ, Lin CY, Kagey MH, Rahl PB, Lee TI, Young RA (2013b) Master transcription factors and mediator establish super-enhancers at key cell identity genes. Cell.

Wickham H (2016) ggplot2: Elegant Graphics for Data Analysis. Springer-Verlag New York.

Winick-Ng W et al. (2021) Cell-type specialization is encoded by specific chromatin topologies. Nature 599:684–691 Available at: http://www.ncbi.nlm.nih.gov/pubmed/34789882.

Wong MM, Byun JS, Sacta M, Jin Q, Baek SJ, Gardner K (2014) Promoter-bound p300 complexes facilitate post-mitotic transmission of transcriptional memory. PLoS One 9 Available at: https://pubmed.ncbi.nlm.nih.gov/24945803/ [Accessed April 4, 2026].

Xiong X, James BT, Boix CA, Park YP, Galani K, Victor MB, Sun N, Hou L, Ho LL, Mantero J, Scannail AN, Dileep V, Dong W, Mathys H, Bennett DA, Tsai LH, Kellis M (2023) Epigenomic dissection of Alzheimer’s disease pinpoints causal variants and reveals epigenome erosion. Cell 186:4422–4437.e21 Available at: https://pubmed.ncbi.nlm.nih.gov/37774680/ [Accessed April 9, 2026].

Yap E-LL, Greenberg ME (2018) Activity-Regulated Transcription: Bridging the Gap between Neural Activity and Behavior. Neuron 100:330–348 Available at: https://linkinghub.elsevier.com/retrieve/pii/S0896627318309012.

Zhang L, Sheng C, Zhou F, Zhu K, Wang S, Liu Q, Yuan M, Xu Z, Liu Y, Lu J, Liu J, Zhou L, Wang X (2021a) CBP/p300 HAT maintains the gene network critical for β cell identity and functional maturity. Cell Death Dis 12:476 Available at: https://pmc.ncbi.nlm.nih.gov/articles/PMC8116341/ [Accessed July 14, 2026].

Zhang MJ, Pisco AO, Darmanis S, Zou J (2021b) Mouse aging cell atlas analysis reveals global and cell type-specific aging signatures. Elife 10.

Zhang Y, Liu T, Meyer CA, Eeckhoute J, Johnson DS, Bernstein BE, Nussbaum C, Myers RM, Brown M, Li W, Shirley XS (2008) Model-based analysis of ChIP-Seq (MACS). Genome Biol.

Zhu LJ, Gazin C, Lawson ND, Pagès H, Lin SM, Lapointe DS, Green MR (2010) ChIPpeakAnno: A Bioconductor package to annotate ChIP-seq and ChIP-chip data. BMC Bioinformatics.

