## Supplementary material for "KAT3 Shuttling Between Neuronal Identity and Activity-Dependent Plasticity Programs Drives Large-Scale Chromatin Remodeling": Supp Figures S1 to S5

### Supplementary Figures and Figure Legends

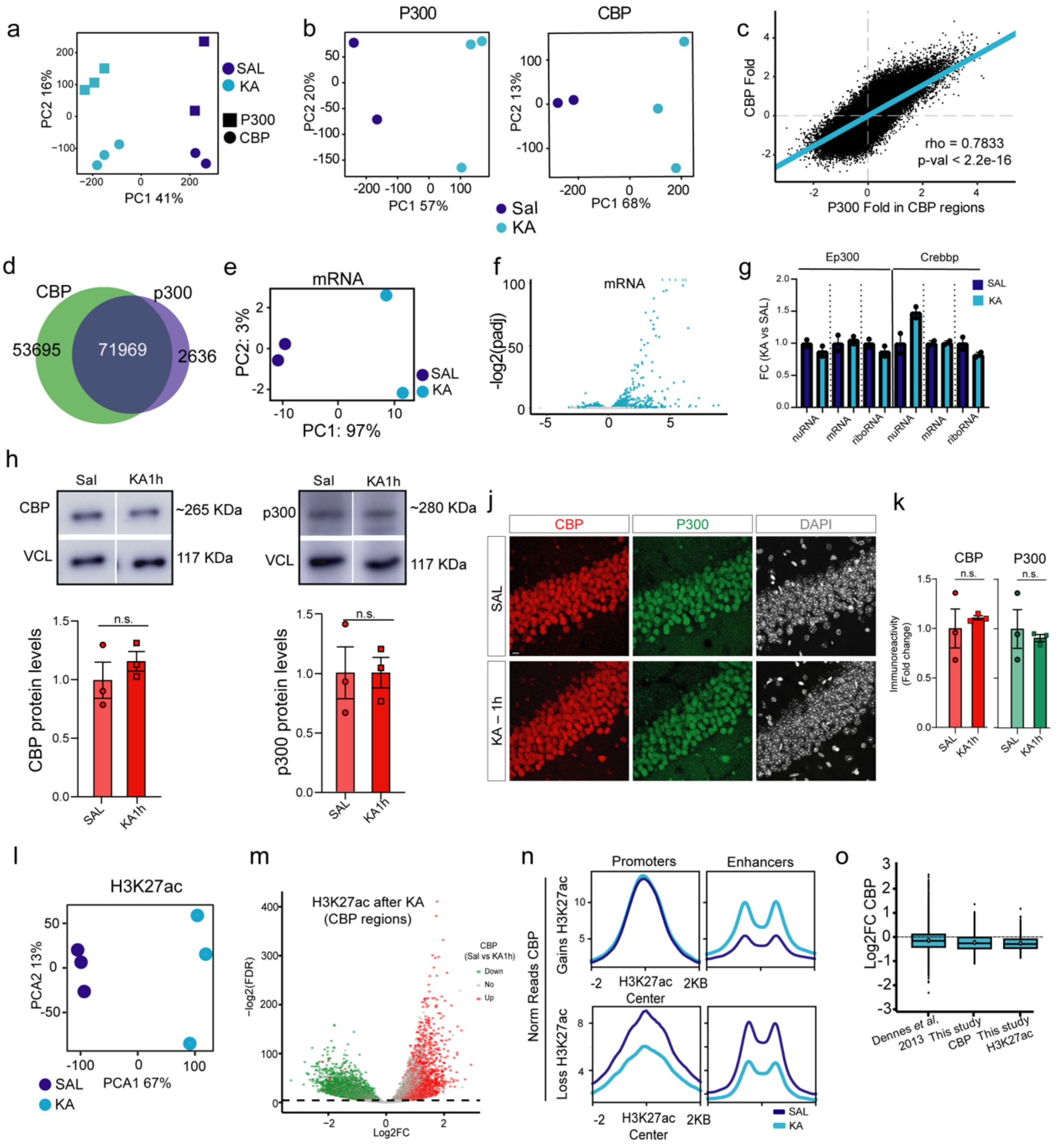

**Supplementary Figure 1 related to Figure 1. CBP/p300 redistribution alters H3K27 acetylation and chromatin accessibility. a.** Combined PCA for p300 and CBP ChIP-seq samples, showing that the major source of variation is activation state. **b.** Principal component analysis (PCA) of p300 and CBP ChIP-seq datasets analyzed separately following KA-induced activation. **c.** Spearman correlation plot comparing CBP and p300 signal intensities at CBP-bound regions. **d.** Venn diagram illustrating the extensive overlap between CBP- and p300-bound regions. **e.** PCA for mRNA-seq samples. **f.** Volcano plot showing differentially expressed mRNAs following KA treatment. **g.** CBP RNA expression levels across three datasets: bulk mRNA from this study, nuclear RNA (nuRNA), and ribosome-bound RNA (riboRNA) from Fernandez-Albert et al. (2019). **h-i.** Western blot (WB) analysis of CBP (h) or p300 (i) in saline and 1 h post-KA samples. Top: representative blot; bottom: quantification. **j.** IHC images of CA1 neurons showing CBP and p300 signals in saline and 1 h post-KA samples. Scale bar = 20 µm. **k**. Quantification of CBP and p300 immunoreactivity in panel j. **l.** PCA for H3K27ac ChIP-seq samples. n = 3 mice per condition. **m.** Volcano plot showing regions with differential H3K27ac following KA treatment. **n.** CBP occupancy at regions showing differential H3K27ac. **o.** Quantification of CBP levels at super-enhancers identified in mouse and human neurons.

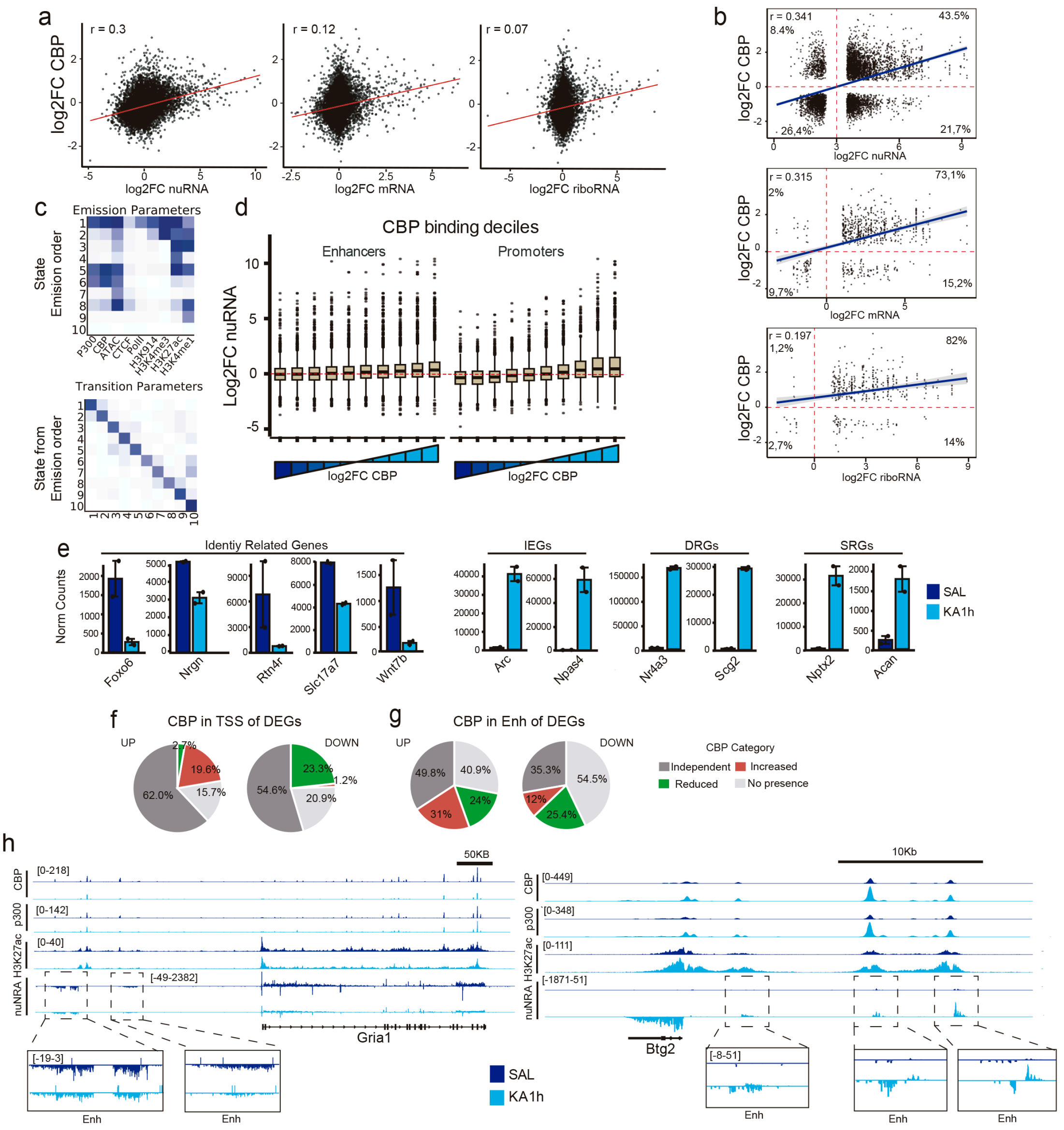


**Supplementary Figure S2 related to Figure 2. Transcriptomic impact of CBP dynamics.** **a.** Correlation between activity-induced changes in CBP binding and changes in nuRNA, mRNA, and ribosome-associated RNA abundance. **b.** Correlation of activity-induced changes in CBP binding and nuRNA-seq, mRNA-seq and riboRNA-seq after restricting the analysis to significantly changed transcripts and CBP peaks. **c.** ChromHMM heatmap showing the chromatin states used to define transcriptionally active regions based on CBP, H3K27ac, and RNA Pol II enrichment. **d.** CBP/p300 peaks separated into TSS-associated and non-TSS regions (putative enhancers) and stratified into deciles according to activity-induced changes in occupancy; corresponding nuRNA changes associated with each group are shown. **e.** Bar plots showing normalized expression levels in saline and KA 1 h conditions for selected genes. nuRNA and riboRNA data are from Fernandez-Albert et al. (2019). **f-g.** Pie charts showing the distribution of CBP peak categories at the TSSs (a) or putative enhancers (b) of nuRNA-seq DEGs. **h.** Genome browser snapshots of representative loci illustrating activity-dependent changes in CBP/p300 occupancy, H3K27ac, and enhancer transcription at putative enhancer regions.

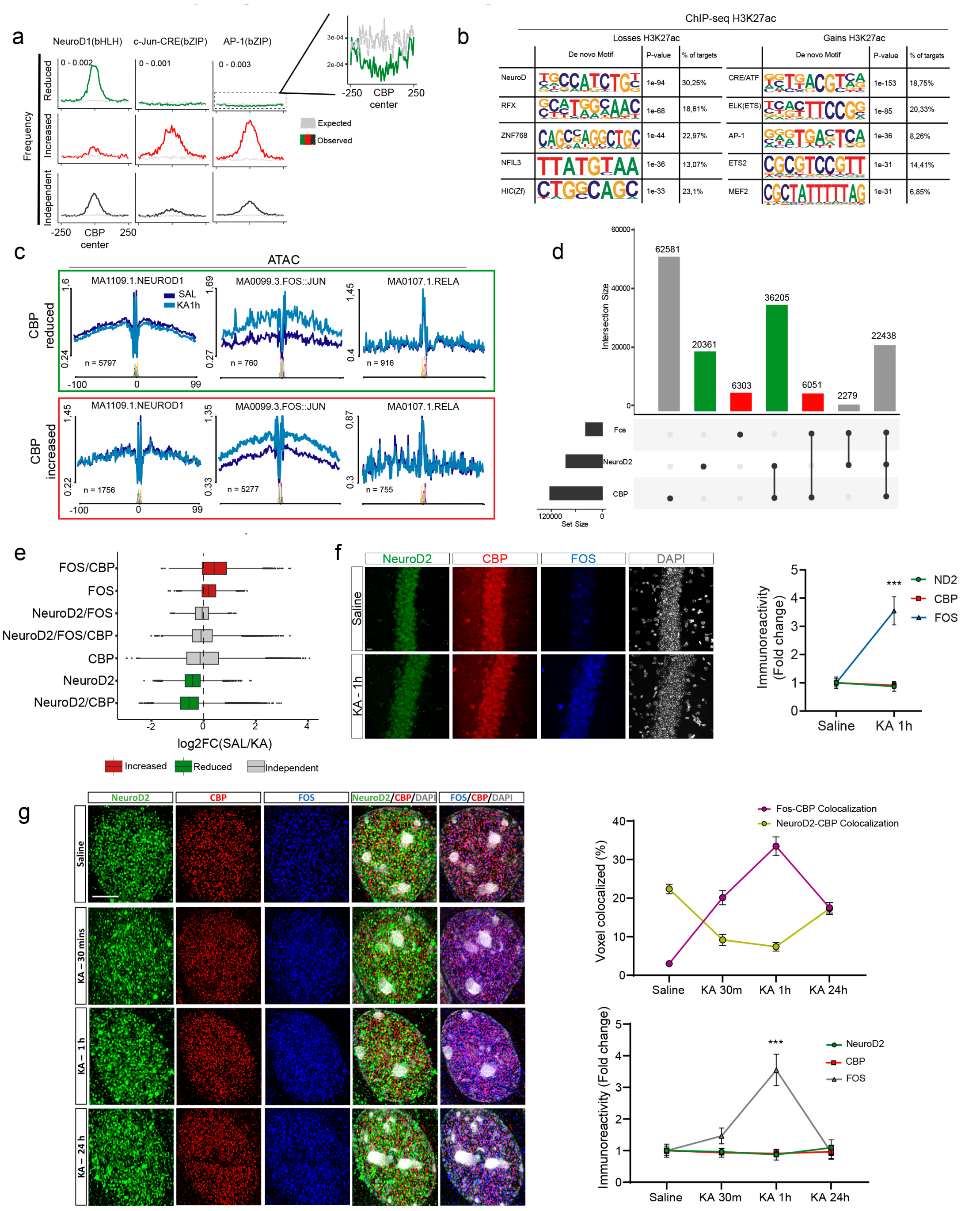


**Supplementary Figure S3 related to Figure 3. Activity-induced AP-1 recruitment redistributes CBP/p300 away from NeuroD-associated loci. a.** Frequency and positional distribution of NeuroD1, CRE, and AP-1 motifs across the three CBP-bound categories. Activity-reduced regions showed a modest depletion of CRE and AP-1 motifs relative to background. **b.** Motif enrichment analysis at regions showing activity-dependent gains or losses of H3K27ac. **c.** Digital footprinting profiles for selected transcription factors at activity-reduced and activity-increased CBP-bound regions. **d.** UpSet plot showing intersections among CBP-, NeuroD2-, and FOS-bound regions. **e.** CBP occupancy under saline and 1 h post-KA conditions across the different NeuroD2/FOS co-binding categories. **f.** Representative confocal images (left) and quantification (right) of nuclear CBP, NeuroD2, and FOS levels before and 1 h after KA treatment. n = 3 mice per condition. Scale bar: 20 µm. **g.** Left: representative super-resolution and deconvolved images showing CBP colocalization with NeuroD2 and FOS within CA1 hippocampal neuron nuclei at different time points after KA treatment. Right: quantification of CBP–NeuroD2 and CBP–FOS voxel co-labeling (top) together with the corresponding immunoreactivity signals (bottom). n = 10 cells/3 mice per condition. Scale bar: 5 µm.

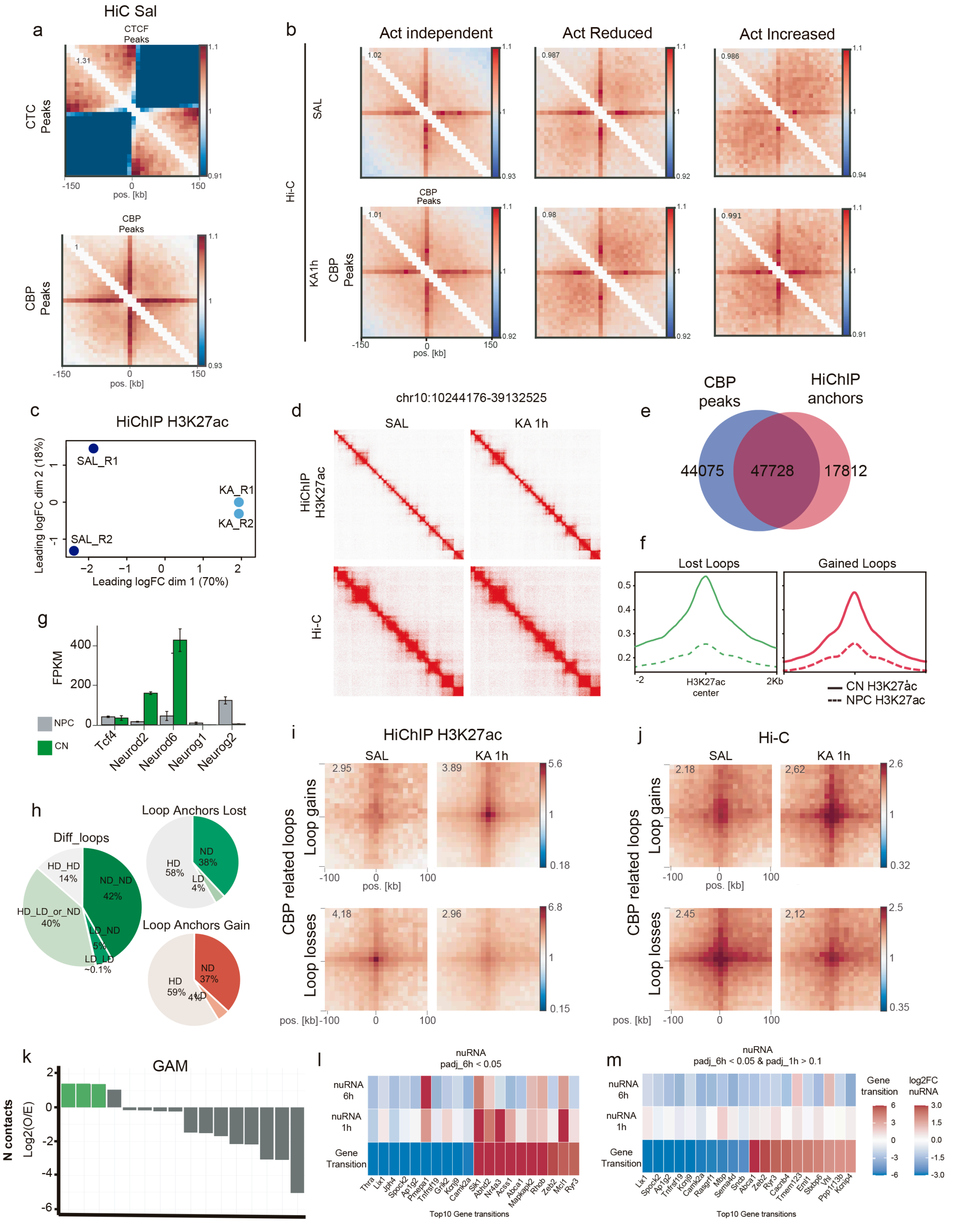


**Supplementary Figure S4 related to Figure 4. Differential loop dynamics and CBP redistribution after neuronal stimulation. a.** Aggregate Hi-C contact maps centered on CTCF peaks (top) or CBP-bound regions (bottom) in hippocampal neurons under basal conditions. **b.** Aggregate Hi-C contact maps centered on activity-independent, activity-reduced, and activity-increased CBP-bound regions under saline and 1 h post-KA conditions. **c.** PCA of H3K27ac HiChIP samples. **d.** Juicer heatmaps of a representative genomic region comparing H3K27ac HiChIP and Hi-C contact maps under saline and 1 h post-KA conditions. **e.** Venn diagram showing the overlap between CBP peaks and H3K27ac HiChIP anchors. **f.** H3K27ac signal in neural progenitor cells (NPCs) and cortical neurons (CNs) at anchors of differential H3K27ac HiChIP loops. **g.** RNA-seq expression levels (FPKM) of selected proneural transcription factors, including NeuroD-family members, in NPCs and CNs. **h.** Left: pie chart showing the relationship between differential HiChIP loops and H3K27ac changes at their anchors. Right: distribution of H3K27ac changes at anchors of lost and gained loops. D = differential H3K27ac; ND = no significant differential H3K27ac. **i.** Aggregate H3K27ac HiChIP signal at differential loops intersecting regions showing activity-induced CBP gain or loss. **j.** Aggregate Hi-C signal at the same differential H3K27ac HiChIP loops intersecting regions showing activity-induced CBP gain or loss. **k.** Analysis of the top 10% strongest CA1 neuron interactions detected by Genome Architecture Mapping (GAM), showing the observed-to-expected enrichment of contacts according to the presence of NeuroD2 and/or CBP at interaction anchors. Expected values were obtained by chromosome-constrained randomization of ChIP-seq peaks while preserving their overall genomic distribution. **l.** Individual spatial-state transitions and corresponding nuRNA changes at 1 and 6 h for selected genes significantly differentially expressed 6 h after KA treatment (adjusted $P < 0.05$). **m.** Individual spatial-state transitions for selected delayed-response genes that were significantly differentially expressed at 6 h (adjusted $P < 0.05$) but showed no significant transcriptional change at 1 h (adjusted $P > 0.1$), together with their corresponding nuRNA changes at both time points.

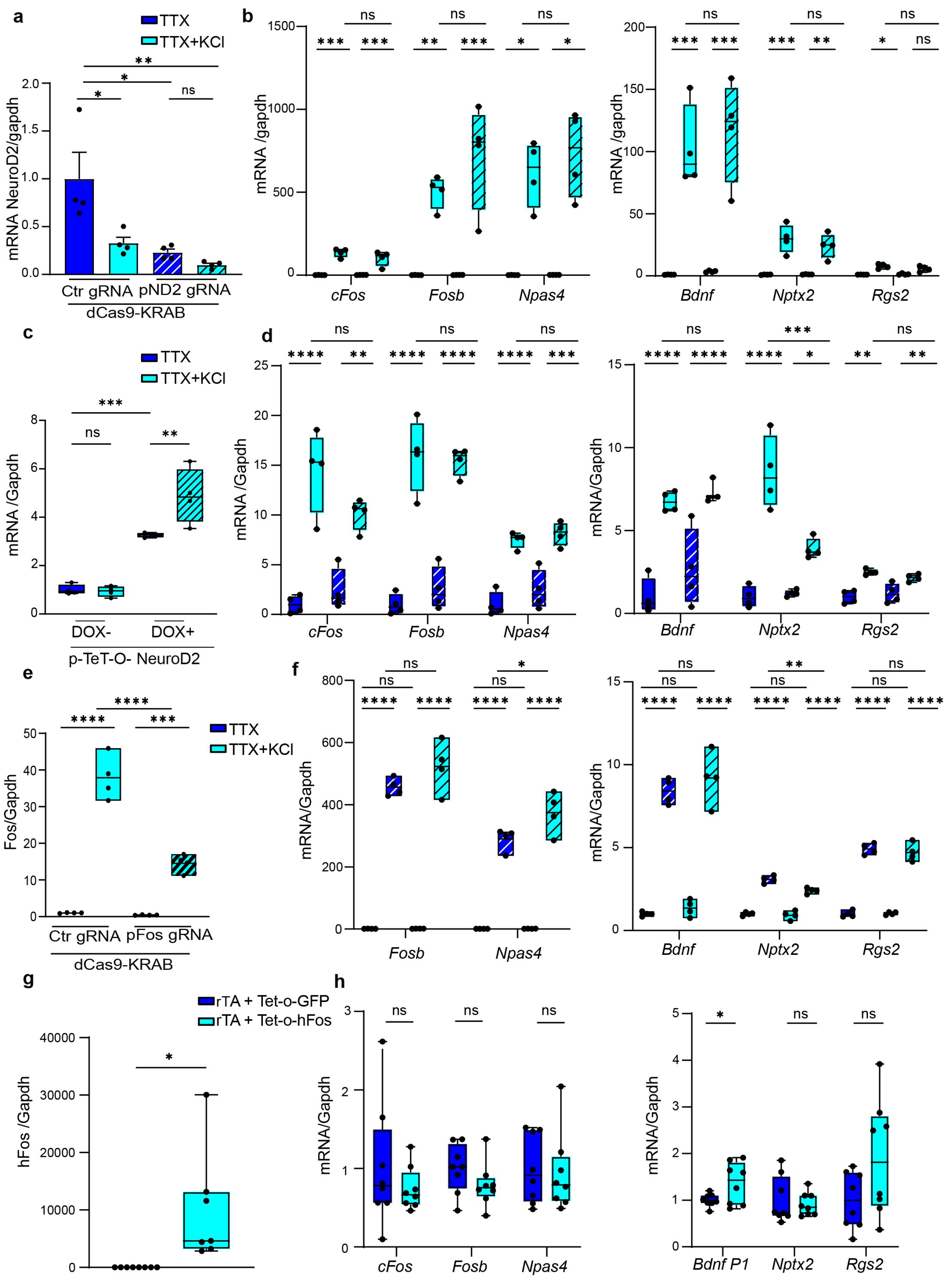


**Supplementary Figure S5 related to Figure 7. Functional loss- and gain-of-function assays. a-b.** NeuroD2 loss-of-function analysis. RT–qPCR confirmed efficient NeuroD2 downregulation (a) and showed no significant effect of NeuroD2 depletion on activity-induced first- and second-wave genes (b). n = 4; mean ± SEM; one-way ANOVA. **c-d.** NeuroD2 gain-of-function analysis. RT–qPCR confirmed efficient NeuroD2 overexpression (c) and showed no significant effect of NeuroD2 upregulation on activity-induced first- and second-wave genes (d). n = 4; mean ± SEM; one-way ANOVA. **e-f.** FOS loss-of-function analysis. RT–qPCR confirmed efficient reduction of Fos expression (e) and showed no significant effect of Fos knockdown on activity-induced first- and second-wave genes (f). n = 4; mean ± SEM; one-way ANOVA. **g-h.** FOS gain-of-function analysis. RT–qPCR confirmed robust induction of FOS expression (g), whereas early- and late-response genes remained largely unaffected (h).
